# Highly multiplexed mRNA quantitation with CRISPR-Cas13

**DOI:** 10.1101/2023.08.16.553527

**Authors:** Brian Kang, Jiayu Zhang, Michael P. Schwoerer, Amy N. Nelson, Emily Schoeman, Andrew Guo, Alexander Ploss, Cameron Myhrvold

## Abstract

RNA quantitation tools are often either high-throughput or cost-effective, but rarely are they both. Existing methods can profile the transcriptome at great expense or are limited to quantifying a handful of genes by labor constraints. A technique that permits more throughput at a reduced cost could enable multi-gene kinetic studies, gene regulatory network analysis, and combinatorial genetic screens. Here, we introduce **q**uantitative **C**ombinatorial **A**rrayed **R**eactions for **M**ultiplexed **E**valuation of **N**ucleic acids (qCARMEN): an RNA quantitation technique which leverages the programmable RNA-targeting capabilities of CRISPR-Cas13 to address this challenge by quantifying over 4,500 gene-sample pairs in a single experiment. Using qCARMEN, we studied the response profiles of interferon-stimulated genes (ISGs) during interferon (IFN) stimulation and flavivirus infection. Additionally, we observed isoform switching kinetics during epithelial-mesenchymal transition. qCARMEN is a simple and inexpensive technique that greatly enhances the scalability of RNA quantitation for novel applications with performance similar to gold-standard methods.

## Introduction

Gene expression levels are indispensable for understanding the functional states of biological systems and can explain the role of certain genes across environmental conditions. Frequently, the expression levels of dozens, hundreds, or even thousands of genes change in response to environmental stimuli or genetic perturbations. The two standard methods for quantifying gene expression changes are quantitative reverse transcription polymerase chain reaction (RT-qPCR) and RNA sequencing (RNA-seq)^1^. While both methods are capable of probing differential gene expression, neither method is effective for quantifying dozens of genes across hundreds or thousands of samples. RNA-seq is too expensive for such high-throughput analyses, costing hundreds of dollars per sample. Conversely, RT-qPCR is too labor-intensive for highly multiplexed studies and consumes large volumes of often precious samples when used to detect more than 4 targets. As a result, gene expression analyses typically involve an initial screen using RNA-seq and subsequent profiling of a few key genes across treatment conditions using RT-qPCR. A scalable, cost-effective method for quantifying RNA would enable gene regulation analysis at the pathway level with high temporal resolution, in response to multiple genetic or chemical perturbations.

An emerging paradigm of nucleic acid detection leverages the sequence-specific *cis-* and collateral *trans-*cleavage activities of type V and VI CRISPR-Cas systems^2–6^. Recent work in viral diagnostics has shown that Cas13a, an RNA-guided RNA nuclease, can be used in conjunction with RNA cleavage reporters to both detect and quantify viral targets such as SARS-CoV-2 and influenza A virus^7–9^. Cas13-based nucleic acid assays are sensitive, detecting targets at concentrations as low as one copy per microliter, and are more specific than nucleic acid hybridization. However, Cas13 detection reactions have continuous kinetics, unlike RT-qPCR, whose discrete cycles allow for differential cycle threshold (ΔCt) analysis. Thus, it has been challenging to infer input RNA concentrations from Cas13 detection kinetics to quantify gene expression in a highly multiplexed fashion.

To address this problem, we have created a Cas13-based platform called qCARMEN that enables high-throughput, multiplexed quantification of changes in host gene expression at a low cost and with minimal hands-on time. The qCARMEN platform uses a system of microfluidics, Cas13 detection reactions, and mathematical models to quantify target RNA concentrations in samples. We demonstrate the utility of the qCARMEN platform for characterizing gene expression kinetics in response to type I and III IFN stimulation, infections with a broad array of genetically diverse flaviviruses, and for studying isoform switching during epithelial-mesenchymal transition in breast cancer cells lines. Furthermore, the gene expression changes calculated using qCARMEN have a high degree of correlation with existing gold-standard methods.

## Results

### Overview of the qCARMEN workflow

The qCARMEN platform enables high-throughput, multiplexed RNA quantitation by parallelizing Cas13-based nucleic acid detection reactions and processing fluorescent readouts using a mathematical model of the reaction kinetics. A typical Cas13 detection reaction leverages the *trans*-cleavage activity of Cas13 to cleave reporter molecules that consist of a fluorophore and quencher separated by a cleavable ssRNA sequence in the presence of target molecules^5^. Prior work has demonstrated that these reactions can sensitively detect viral targets and that the rate of fluorescence saturation correlates with target concentration^5^. However, CRISPR-based nucleic acid quantitation has proven to be challenging due to competition between amplicons and primer dimers during multiplexed amplification and the lack of a robust method for calculating nucleic acid concentrations from fluorescence kinetics **(Extended Data Fig. 1)**. qCARMEN addresses these challenges by integrating a primer dimerization-resistant amplification strategy and reaction kinetics modeling using a system of differential equations.

The entire qCARMEN workflow requires less than two hours of total hands-on time and consists of three primary steps: complementary DNA (cDNA) synthesis, pre-amplification, and microfluidic chip loading **(Fig. 1a)**. After cDNA synthesis, samples are pre-amplified in a multiplexed format using an RNase H2-dependent PCR (rhPCR) protocol to increase assay sensitivity and to incorporate a T7 promoter sequence into target amplicons^10^. When compared to pooled amplification with conventional primer sets, rhPCR allows for significantly higher gene coverage in multiplexed amplification and eliminates primer dimers **(Extended Data Fig. 2)**. PCR products are then loaded onto a microfluidic chip, where samples are combinatorially mixed with assay master mixes consisting of a target-specific CRISPR RNA (crRNA), LwaCas13a, and T7 RNA polymerase^7^. Upon mixing, reactions are incubated for three hours at 37°C with fluorescence measurements taken every five minutes. Raw fluorescence data are fit to a mathematical model to calculate relative target concentrations, which can be used to quantify differential expression across treatment conditions.

**Fig. 1.**
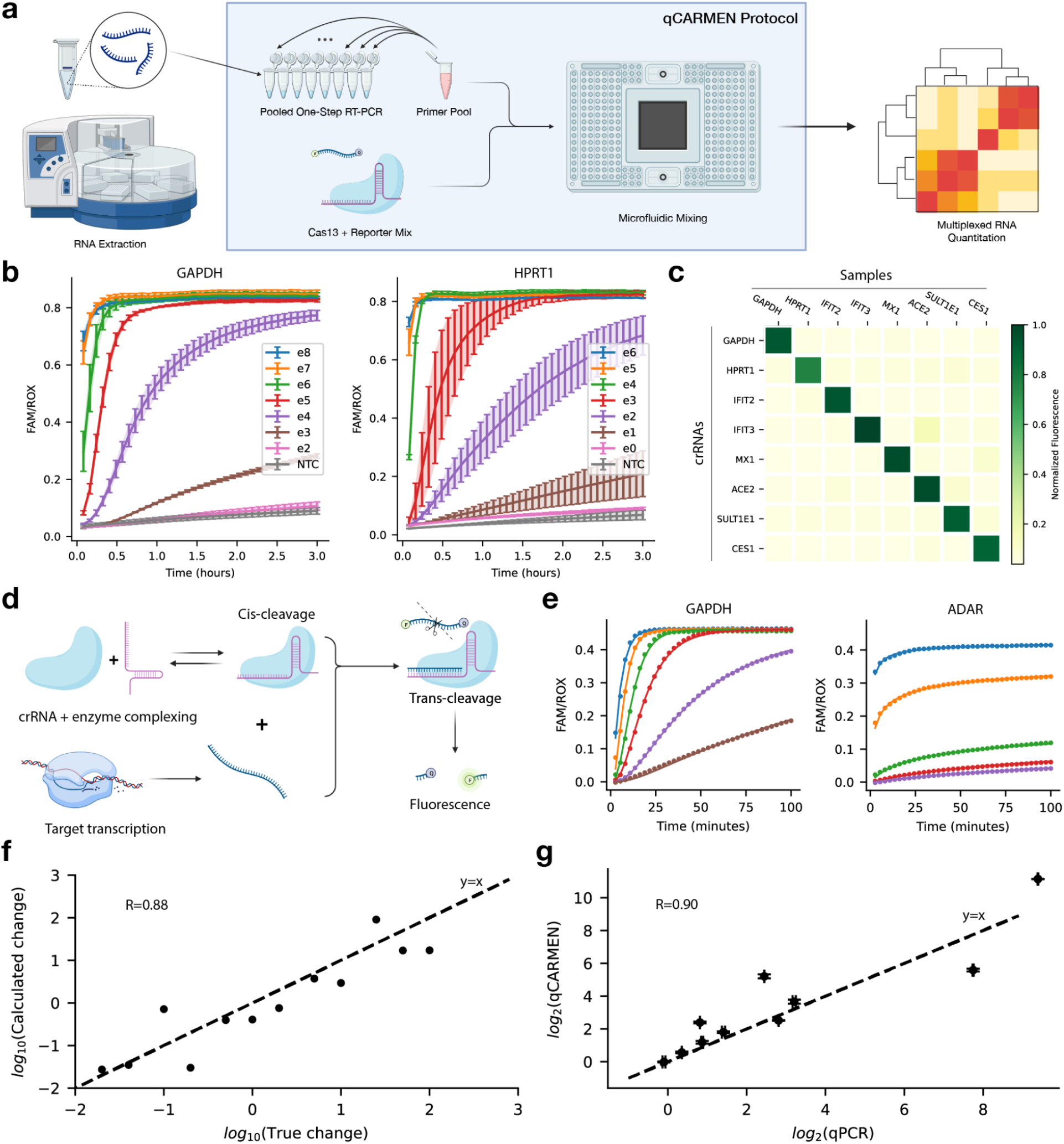
Validating the sensitivity, specificity, and accuracy of qCARMEN. **a**, Following an RNA extraction step, samples are amplified using commercially available rhPCR primers and combined with a Cas13 assay mix in a microfluidic chip. Reactions run for three hours and fluorescent signals are recorded for downstream analysis. **b**, Serial dilutions of Huh7.5 cellular RNA extracts show sensitive detection of housekeeping genes as low as 1-10 copies/μL depending on the target of interest. Error bars presented as 95% confidence intervals. **c**, Cas13 reporter assays generate fluorescent signals in the presence of a target that is at least 5-fold greater than the background fluorescence. **d**, Simplified model of the Cas13 detection reaction. **e**, The mathematical model generates precise fits to real-world fluorescence data. **f**, Calculated fold-changes for serially diluted synthetic HPRT1 RNA targets plotted against dilution factors. **g**, Huh7 cells were infected with YFV-17D at a MOI of 0.05 and fold-changes in expression of ten genes at 2 dpi were calculated using RT-qPCR and qCARMEN with considerable agreement between the two methods. Error bars presented as SEM (n=3).

### Cas13-based transcript detection is highly sensitive and specific

To validate the sensitivity of qCARMEN, we generated synthetic transcripts of the housekeeping genes glyceraldehyde 3-phosphate dehydrogenase (GAPDH) and hypoxanthine phosphoribosyltransferase 1 (HPRT1) via *in vitro* transcription (IVT). Synthetic housekeeping gene samples were serially diluted from an initial concentration of 10^12^ copies/μL by a factor of ten to generate transcript samples with concentrations ranging from 1 to 10^8^ copies/μL. To establish the limit of detection (LOD) of the qCARMEN reaction for each housekeeping gene, we used a no template control (NTC) consisting only of nuclease-free water as the input for two reactions separately aiming to detect either GAPDH or HPRT1. The serial dilutions were used as inputs for qCARMEN reactions detecting GAPDH and HPRT1, and fluorescence outputs were collected. Fluorescence data were compared to the NTC for each gene to establish a comparative LOD. Based on the NTC outputs, we determined limits of detection for both GAPDH and HPRT1 to be approximately 100-1,000 copies/μL and 1-10 copies/μL, respectively **(Fig. 1b)**.

To test the specificity of the qCARMEN workflow, eight genes were individually amplified in total RNA extracted from human Huh7.5 hepatoma cells^11^. Each of the eight PCR products was then combined with eight unique assay mixes each containing a crRNA targeting one of the eight target genes for a total of 64 assay-sample mix combinations. Cas13-mediated detection of target genes was highly specific, generating a positive signal approximately 15 times above background (**Fig. 1c**).

### Modeling the Cas13 detection reaction for effective transcript quantitation

To quantify changes in gene expression using fluorescence data from Cas13 detection reactions, fluorescence curves need to be converted into scalar values that correlate with input RNA concentrations. Prior work has inferred input RNA concentrations by calculating half-maximal inhibitory concentration (IC_50_) values from fluorescence kinetic curves; however, this requires saturating reaction kinetics and therefore severely limits the dynamic range of gene expression quantitation^7^. To circumvent this limitation, we modeled the Cas13 detection system using a system of differential equations representing individual reaction components including transcription, *cis* cleavage, and *trans* cleavage **(Supplementary Note 1, 2)**. Parameters generated for the model were largely consistent across different targets, exhibited linearity in predicted concentration trends, and were able to fit distinct types of fluorescence kinetics **(Extended Data Fig. 3)**. By using our model, we were able to create precise fits to our fluorescence data and generate a single concentration-associated parameter for each curve **(Fig. 1d, 1e)**.

With the derived model parameters, we aimed to accurately and precisely quantify changes in gene expression. To validate our model, we first generated dilutions of a synthetic HPRT1 RNA target and used our qCARMEN assay to generate fluorescence curves for each dilution. Then, by fitting our model to the fluorescence data, we calculated changes in input RNA levels across these dilutions and found that log_10_-transformed calculated and true fold-change values were approximately one-to-one with each other **(Fig. 1f)**. Next, we validated the model in a cellular context by infecting Huh7 cells with the yellow fever virus vaccine strain 17D (YFV17D) and observing changes in expression across 10 genes selected from an RNA-Seq analysis of Huh7 cells harboring subgenomic replicons (SGRs) of YFV17D or the virulent YFV-Asibi parental strain (**Extended Data Fig. 4**). After extracting RNA from mock and infected cells, we used both RT-qPCR and qCARMEN to independently determine expression changes. By plotting the log_2_ fold-changes from the two methods against each other, we found high one-to-one linearity between RT-qPCR and qCARMEN results with a correlation coefficient of 0.90 (**Fig. 1g**). The results from these two experiments demonstrate that qCARMEN can accurately quantify gene expression changes in consensus with qPCR.

### Analysis of IFN response kinetics using the qCARMEN ISG panel

The IFN response plays a key role in fighting infection, and studying ISGs can provide mechanistic insights into pathogen defense pathways^12^. IFN-mediated cellular responses to infections have been studied in great detail and involve hundreds of ISGs, thus lending themselves as a test case for qCARMEN. Examples of well-characterized ISGs include the oligoadenylate synthetase (OAS) family, which is responsible for detecting foreign RNA, and the IFN-induced transmembrane (IFITM) proteins, which typically inhibit viral endocytic fusion events^13^. Thus, we sought to design a qCARMEN panel covering approximately two dozen commonly induced ISGs^13–15^.

We first applied the ISG panel towards studying differences in ISG dysregulation in response to type I and III IFN stimulation^12^. While both type I and III IFNs can induce antiviral states, they generate distinct ISG response profiles (Mesev et al., in revision). Type I IFN stimulation typically triggers a fast, but robust ISG response whereas type III IFN stimulation induces weaker, but longer lasting responses. We stimulated human bone osteosarcoma epithelial (U2OS) cells, which bear both type I and III interferon receptors, with IFNs β, α2, and λ3 (referred to as IFNs β, α, and λ hereafter) at doses that we previously determined to generate strong ISG responses (Mesev et al., in revision). IFN-stimulated and mock cells were taken down over the course of 72 hours, after which RNA was extracted and quantified using qCARMEN for all ISGs that exhibited baseline expression in mock cells.

By using qCARMEN to quantify changes in gene expression in IFN-stimulated cells relative to mock samples, we found that most ISGs were upregulated across type I and III IFN-stimulated cells, as expected based on prior results (Mesev et al., in revision) and publicly available data on IFN-mediated ISG dysregulation (**Fig. 2b**)^16^. Additionally, type I and III IFN stimulation generally induced the same direction of ISG dysregulation for most genes in the panel after 24 hours (**Fig. 2c**). However, for two ISGs—moloney leukemia virus 10 (MOV10) and single-stranded DNA binding protein 3 (SSBP3)—we observed different directionality in ISG changes across the IFN subtypes.

**Fig. 2.**
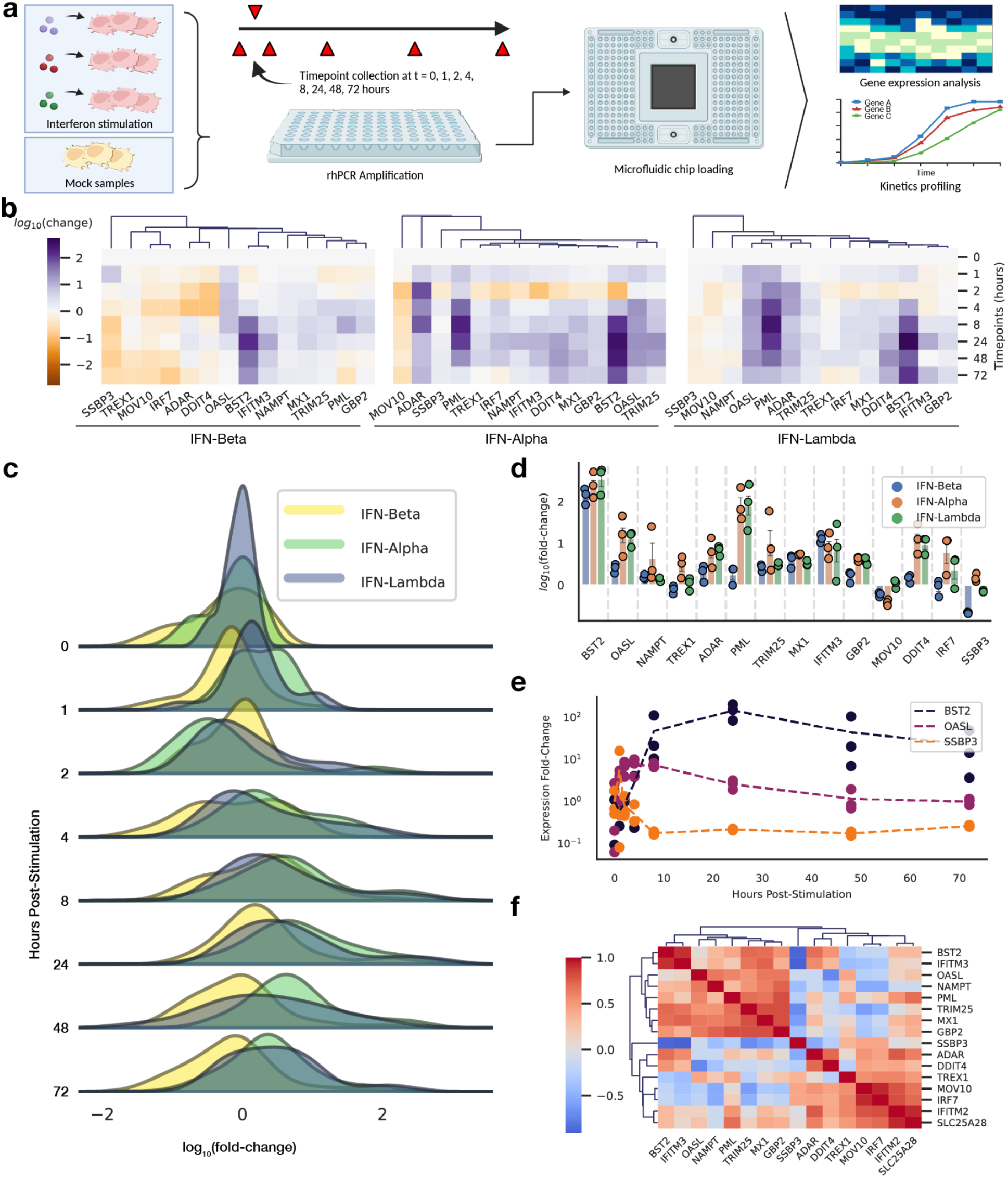
Development of an interferon-stimulated gene (ISG) panel. **a**, U2OS cells were stimulated with IFNs β, α, and λ to dysregulate ISG expression. Timepoint samples were collected over the course of 72 hours for stimulated and naïve cells for downstream analysis with qCARMEN. **b**, Gene expression changes of ISGs after IFN stimulation (120 IU/mL of IFN-β, 800 IU/mL of IFN-α, 40 ng/mL of IFN-λ) across all eight timepoints for type I and III IFN experiments. **c**, Ridge-line plots showing overall changes in kinetics across stimulated ISGs. **d**, log_2_ fold-changes in ISG expression levels after 24 hours. Error bars presented as SEM (n=3). **e**, Tetherin (BST2), OASL, and SSBP3 demonstrate differential kinetics all within IFN β-stimulated cells. **f**, Pairwise Spearman correlations for IFN β-stimulated genes; simple average used for linkage to calculate clusters with euclidean distance metric.

Plotting the distribution of fold-change values at each time point shows the differences in kinetics and response strength for each IFN (**Fig. 2d**). Stimulation with IFN-β, a type I IFN, generated a rapid ISG response that peaked at 4 hours. Overall, IFN-λ induced a slower ISG response that peaked at 24 hours—much later than with IFN-β—which is consistent with previous observations^17–21^.

The data from the qCARMEN panel also allowed for a more in-depth kinetic analysis of specific genes. As previously mentioned, type I IFN responses can be characterized by swift and marked changes in expression^18,21^. However, this generalization does not necessarily apply to all downstream ISGs^18^. By specifically looking at the kinetic profiles of bone marrow stromal cell antigen 2 (BST2), 2’-5’-oligoadenylate synthetase-like (OASL), and SSBP3 during IFN-β stimulation, we observed distinct examples of up-regulation, down-regulation, and delayed dysregulation all in response to stimulation by the same type I IFN (**Fig. 2e**).

The throughput and multiplexing capabilities of qCARMEN also allowed us to calculate pairwise correlations across genes to assess co-regulation within different interferon responses. As expected, when calculating Spearman correlations for genes responding to IFN stimulation, we found that most ISGs were positively correlated with one another **(Fig. 2f)**. However, some genes—SSBP3, adenosine deaminase acting on RNA (ADAR), DNA damage inducible transcript 4 (DDIT4), MOV10, and interferon regulatory factor 7 (IRF7)—exhibited negative correlations in cells stimulated with IFN-β. Similarly, IFN-α and IFN-λ stimulation resulted in high positive correlations for most ISGs, though ADAR and SSBP3 were also negatively correlated in the IFN-α and λ datasets **(Extended Data Fig. 5)**.

### Application of ISG panel towards characterizing host responses to flavivirus infections

Flaviviruses are a family of positive-sense, single-stranded RNA viruses that cause infections in over 500 million people annually^22^. Within this family of viruses, infections with different viruses can cause vastly different clinical outcomes. To demonstrate the broad applicability of our ISG panel and to characterize differences in host cell intrinsic innate immune responses, we infected Huh7 hepatoma cells with an array of flaviviruses. Huh7 cells were infected with dengue virus (DENV) serotypes 1-4, Langat virus (LGTV), Usutu virus (USUV), Zika virus (ZIKV), hepatitis C virus (HCV), or YFV-17D **(Fig. 3a)**. The frequencies of viral antigen-bearing cells were determined via flow cytometry and cells were lysed to extract total RNA in bulk from infected and mock cells two to five days post-infection (dpi) **(Extended Data Fig. 6)**. RNA from mock Huh7 cells was used to generate a dilution series for quantitation. Additionally, to ensure that dysregulated genes that may not be present in high concentrations in mock cells were represented in our dilution series, we used total RNA from DENV-4-infected cells to generate another dilution series more representative of the infected state.

**Fig. 3.**
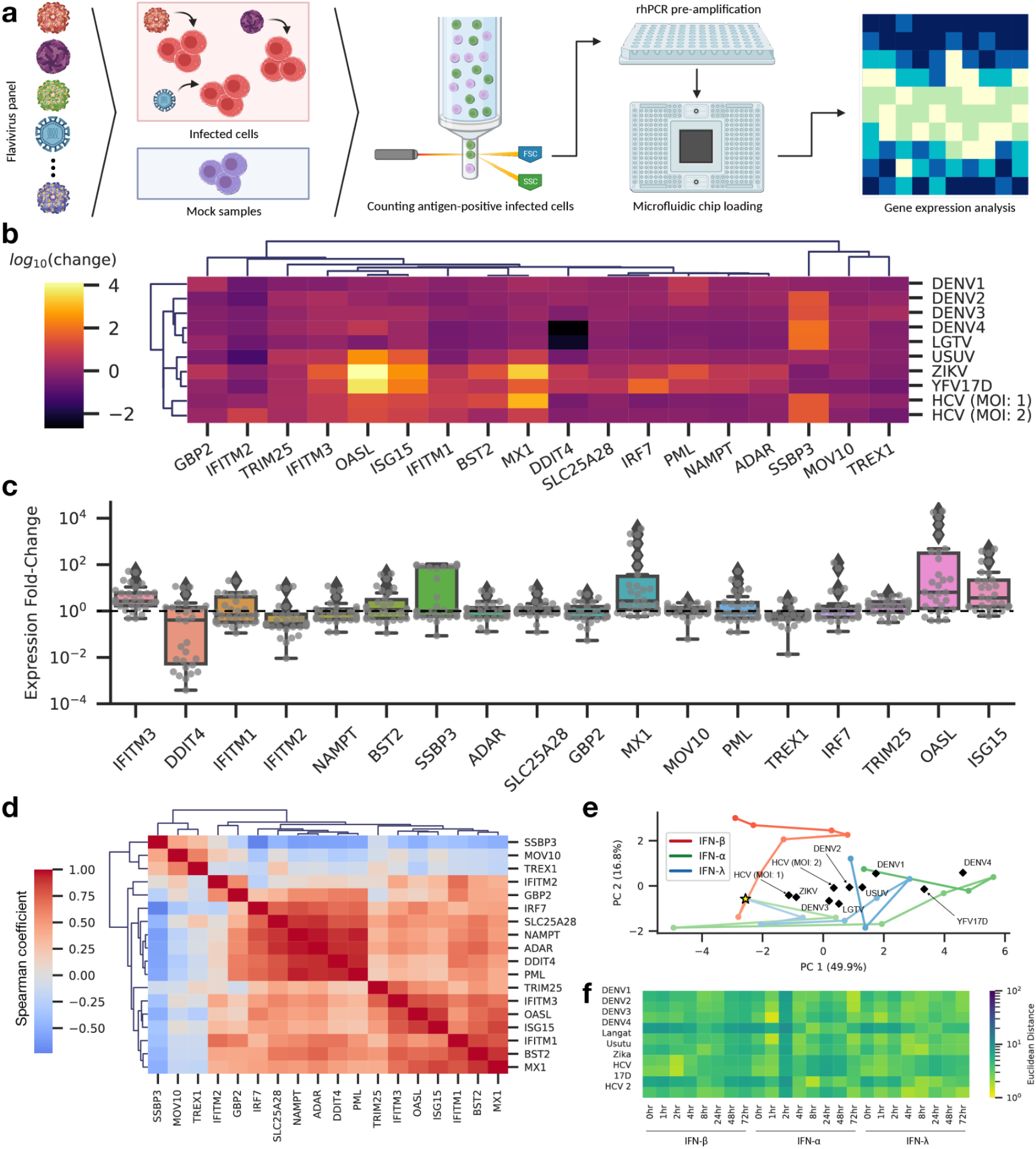
Application of the ISG panel towards understanding flavivirus infections. **a**, Huh7 cells were infected with nine flaviviruses. Total RNA extracted from infected Huh7 cells was used as input for the qCARMEN ISG panel. **b**, Calculated fold-changes in expression for ISG panel genes across infected cells compared to naive cells. **c**, The distribution of fold-changes for each gene across viral infections and replicates. Boxplot whiskers extend to 1.5 times the interquartile range; outliers represented with diamond markers. **d**, Pairwise Spearman correlations of dysregulated ISGs in response to flavivirus infection. **e**, Gene expression vectors at each timepoint for all three IFNs plotted along two principal components to illustrate the trajectory of expression changes across core ISGs. PCA were transformations performed on flavivirus ISG signatures to overlay the expression profiles on top of IFN stimulation trajectories. Gold star represents 0-hour timepoint. Light to dark: 1, 2, 4, 8, 24, 48, 72-hour timepoints. **f**, Euclidean distance between flavivirus PCA projections and IFN timepoint vectors to characterize flavivirus ISG changes in the context of IFN responses.

The ISG panel showed distinct ISG responses across the different flavivirus infections both in the magnitude and direction of changes as well as the sets of dysregulated genes (**Fig. 3b**). By looking at the distribution of calculated fold-changes across viruses for each gene, we can see that five ISGs in particular—DDIT4, SSBP3, myxoma resistance protein 1 (MX1), OASL, and interferon-stimulated gene 15 (ISG15)—exhibited the greatest variance across the array of flaviviruses (**Fig. 3c**). The expression of DDIT4 decreased in response to most virus infections, particularly for HCV and ZIKV. The latter four highly dysregulated ISGs generally demonstrated an increase in expression with up to four orders of magnitude in variation across viruses. The remaining ISGs in the panel deviated minimally from baseline expression levels for the majority of viruses.

To identify key genes relevant in the immune response to flavivirus infections, we determined Spearman correlations between genes based on the gene signature vectors generated for each infection. The Spearman correlation matrix shows two primary clusters of positively correlated genes. Notably, SSBP3 exhibited strong inverse correlations with the other ISGs on the panel except for MOV10 and three prime repair exonuclease 1 (TREX1). Additionally, SSBP3 and MOV10 both exhibited similar negative correlations with other ISGs when stimulated with type I IFNs β and α **(Fig. 2d** and **Extended Data Fig. 5a)**. The role that SSBP3 plays in the antiviral response is largely elusive; however, it is known that SSBP3 is a transcriptional regulator involved in interfering with viral translation. One possible explanation for the inverse correlation is that SSBP3 plays a role in regulating the IFN response by suppressing ISG expression to avoid an overly extended immune response. Since MOV10 and TREX1 do not exhibit significant changes in expression in response to any of the viral infections, it is logical that SSBP3 is not inversely correlated with these two ISGs (**Fig. 3c**).

Lastly, we sought to compare the ISG responses for each flavivirus infection to the ISG profiles induced by type I and III IFN stimulation. To do so, we performed principal component analysis (PCA) on the gene expression signatures for each IFN and timepoint. Since the timepoints at which infected cells were taken down varied across viruses, incorporating gene expression data from each timepoint allowed us to include a temporal component in our characterization of each flavivirus. Plotting gene expression vectors from each IFN timepoint allowed us to first observe overlapping trajectories in the progression of the ISG responses (**Fig. 3E**). IFNs α and λ exhibited very similar trajectories during the first four hours post-infection, diverged at the 8-hour timepoint, and eventually converged after 72 hours. IFN-β exhibited a completely different trajectory altogether. The host responses of 8/10 viruses were far from the unperturbed 0-hour timepoint. Calculating the Euclidean distance between flavivirus ISG profiles and IFN stimulation timepoints using generated PCA components provided a quantitative description of the trajectory analysis (**Fig. 3E**). Most viruses used here—DENV-1, DENV-2, DENV-3, DENV-4, LGTV, USUV, and YFV17D—had ISG signatures that most closely resembled IFN α or λ responses during the first 1-4 hours of stimulation. The remaining flaviviruses—HCV and ZIKV—had expression signatures most similar to that of earlier timepoints for IFN β (**Fig. 3F**). Collectively, these data demonstrate that qCARMEN is readily suitable to characterize complex transcriptional changes induced by cytokines and viral infection.

### Quantitation of alternative splicing in breast cancer cells

An additional layer of complexity in host transcriptional changes arises from alternative splicing of mRNAs. While quantifying changes in splice isoforms is possible with existing RT-qPCR techniques, designing primer sets that specifically amplify one particular isoform can be challenging due to primer location and design constraints. Because Cas13-based detection enforces specificity through both primer hybridization as well as crRNA-mediated reporter cleavage, qCARMEN stands to offer an effective alternative to RT-qPCR for measuring changes in expression not only across genes, but also across distinct splice isoforms.

To validate qCARMEN for isoform quantitation, we studied isoform switching events in breast cancer cell lines during epithelial-mesenchymal transition (EMT). Studies indicate that EMT plays a meaningful role in tumor progression due to the loss of cell adhesiveness and activation of motility during EMT^23,24^. Additionally, alternative splicing has also been shown to occur during EMT in numerous cancer cell lines^25–28^. EMT can easily be induced in such cell lines by treatment with recombinant transforming growth factor β (TGF-β), allowing us to generate samples with differentially expressed splice isoforms^29^. We designed a qCARMEN panel to observe alternative splicing for CD44, fibroblast growth factor (FGFR1), mothers against decapentaplegic 2 (SMAD2), and exocyst complex component 7 (EXOC7)—genes that have been previously characterized in the context of alternative splicing during EMT. We treated MCF7 and T47D breast cancer cell lines with recombinant human TGF-β and collected samples over the course of 72 hours. Extracted RNA used as input for the qCARMEN workflow.

To first confirm that Cas13 is capable of distinguishing between splice isoforms, we generated synthetic isoforms for FGFR2 and designed crRNAs and primer pairs targeting either the epithelial or the mesenchymal isoform. To assess specificity, isoform-specific assay mixes were combined with both the synthetic epithelial and mesenchymal target mixes to generate a fluorescent readout over the course of three hours. Detection was largely specific for both isoforms with some background detection of the FGFR2 IIIc isoform in the IIIb target mix (**Fig. 4B**). We then created dilutions of each isoform with the other isoform present in solution at a constant concentration across each dilution. Cas13 detection assays generated distinct fluorescence curves for each dilution for both the IIIb and IIIc isoforms of FGFR2, demonstrating that qCARMEN can distinguish different concentrations of splice isoforms and quantify changes in isoform expression (**Fig. 4C**).

**Fig. 4.**
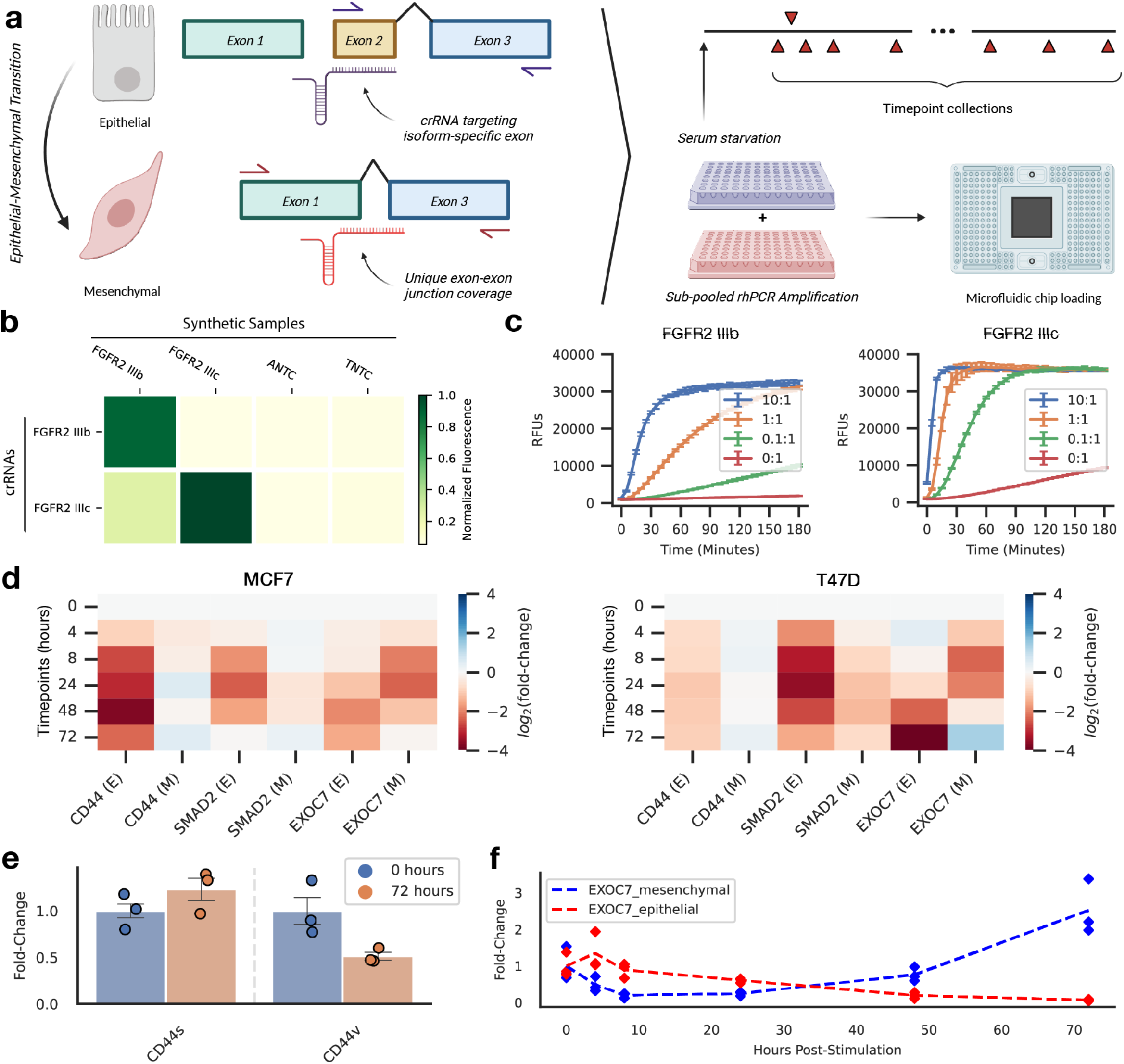
Quantifying changes in dominant splice isoforms during epithelial-mesenchymal transition. **a**, Schematic representation of the experimental design. **b**, Cas13-based detection of synthetic FGFR2 isoforms is largely specific with relatively minimal background compared to positive controls. **c**, Detection of synthetic FGFR2 isoform dilutions. Serial 10X dilutions of IIIb isoform of FGFR2 with IIIc isoform at a constant concentration (left). Serial 10X dilutions of IIIc isoform with IIIb isoform left constant. Error bars presented as 95% confidence intervals. **d**, Calculated changes in expression across downregulated epithelial (E) and upregulated mesenchymal (M) splice isoforms in both MCF7 (left) and T47D (right) cells after treatment with TGF-β. **e**, CD44s (mesenchymal form) increases relative to the initial timepoint after 72 hours; CD44v (epithelial form) decreases approximately 2-fold after 72 hours. Error bars presented as SEM (n=3). **f**, Kinetics of EXOC7 isoforms over a 72-hour timecourse.

Next, we applied qCARMEN towards the analysis of differentially expressed splice isoforms in MCF7 and T47D cells after induction of EMT. The genes in the panel showed the expected increases in expression across the mesenchymal forms relative to the epithelial forms over the course of three days for both cell lines. The changes in gene expression were more noticeable in T47D cells, especially for the mesenchymal splice isoform of EXOC7, which exhibited nearly a 6-fold increase in expression between the 0- and 72-hour timepoints **(Fig. 4D)**. CD44, a cell surface protein known to undergo alternative splicing during EMT, exhibited an increase in its mesenchymal form and a decrease in its epithelial isoform after 72 hours of TGF-β treatment as expected from past studies **(Fig. 4E)**^25^.

In addition to assessing long-term fold-changes in expression, we examined the kinetics of splice isoform expression patterns and identified striking behaviors in the dynamics of the epithelial and mesenchymal isoforms. In particular, during EMT in T47D cells, the epithelial and mesenchymal isoforms of EXOC7 do not exhibit a simplistic, immediate decrease or increase in the dominant isoform **(Fig. 4F)**. During the first two hours of TGF-β stimulation, the mesenchymal EXOC7 isoform decreases relative to the initial timepoint and the epithelial form increases until the 4-hour time point. After reaching these critical points, the epithelial and mesenchymal forms begin to trend towards the expected direction of change, and by the 72-hour time point, we see an increase in the mesenchymal form and a decrease in the epithelial form. Taken together, these results establish proof-of-concept for the utility of qCARMEN to monitor the complex dynamics of different RNA splice forms during EMT transitions.

## Discussion

In summary, we developed and validated qCARMEN: a rapid RNA quantitation tool that is the first of its kind to use CRISPR-Cas13 for high-throughput, multiplexed quantitation of host gene transcripts. To validate qCARMEN experimentally, we designed an ISG-focused panel to survey interferon response profiles in both interferon-stimulated and flavivirus-infected cells. Using our ISG panel, we were able to quantify gene expression changes for nearly two dozen genes across one hundred RNA samples in under two hours of hands-on time. By profiling kinetics across these ISGs, we identified genes such as SSBP3 that were inversely correlated with other ISGs across type I and III interferon stimulation experiments as well as in flavivirus infection samples. In addition to the ISG panel, we validated isoform quantitation using qCARMEN and studied isoform switching during EMT in breast cancer cell lines. The isoform panel verified known changes in splice isoform expression after EMT and uncovered notable kinetic trends in certain transcripts such as EXOC7.

While this study focused primarily on ISG responses and isoform switching during EMT, the qCARMEN panel design process and workflow can be readily applied to any set of genes or RNA samples. We envision the use of qCARMEN across a wide range of applications, especially in the development of machine learning classifiers that require high dimensional gene expression data for accurate and nuanced predictions of clinical outcomes. Moreover, we expect qCARMEN to significantly lower the labor and reagent costs associated with highly multiplexed kinetics studies and gene regulatory network analyses. While it is possible to assay dozens of genes across hundreds of timepoints or treatment conditions with current gold-standard methods, this is rarely done because either the cost or time required is prohibitive. With qCARMEN, we believe that high-throughput, multiplexed studies will become increasingly attractive, which will generate larger datasets that can be used to train machine learning models, perform network analyses, and uncover new insights into pathways of interest.

One limitation of qCARMEN is the current dynamic range as it necessitates an initial experiment to optimize primer concentrations. When designing a new panel from scratch, this will require additional hands-on time to ensure optimal assay performance. However, we find that this is not a major limitation in practice because rhPCR generally allows for effective amplification of most targets with minimal optimization and because qCARMEN does not require a large amount of input material. In future studies, one could use biochemical approaches and thermal regulation to slow the detection reaction to better distinguish between fluorescence outputs of highly concentrated targets and increase the dynamic range.

Current RNA quantitation tools are limited either by throughput or target multiplexing. qCARMEN offers a solution for these limitations by leveraging CRISPR-Cas13 and microfluidics to observe changes in gene expression across dozens of targets and hundreds of samples simultaneously with only two hours of hands-on time. The rhPCR primer and crRNA design process is fully automated for qCARMEN, so new panels targeting other distinct sets of genes can be designed and validated rapidly. In conclusion, the throughput and multiplexing capabilities of qCARMEN can enable a new range of applications of RNA quantitation including disease prognostics and predictive tools by lowering the cost and labor barriers associated with high-throughput RNA quantitation.

## Materials & Methods

### Cell culture

Five cell lines were used in this study: Huh7, Huh7.5 (both kindly provided by Charles M. Rice, The Rockefeller University), U2OS (commercially obtained from the American Type Culture Collection, ATCC, Manassas, Virginia), MCF7, and T47D (both kindly provided by Yibin Kang, Princeton University). Unless otherwise stated, all cells were grown under standard conditions (37°C, 5% (v/v) CO_2_) and cultured in Dulbecco’s Modified Eagle’s Medium (DMEM, Thermo Fisher Scientific, Waltham, MA) containing 10% (v/v) heat-inactivated fetal bovine serum (FBS) and 1% v/v penicillin/streptomycin (Corning Inc., Corning, NY). Upon reaching confluency, cells were trypsinized with 0.05% (v/v) trypsin/EDTA and re-plated on cell culture dishes.

### Design of primers and crRNAs

Primer and crRNA sequences for target transcripts were designed using an automated computational pipeline (https://github.com/Myhrvold-Lab/qCARMEN). The computational pipeline consists of three major components. First, crRNAs are designed for transcripts using ADAPT^30^. Forward and reverse primer sequences are then designed for optimal crRNA designs based on G/C content, desired melting temperature, amplicon length, and the presence of mono- and di-nucleotide repeats. Lastly, T7 promoter sequences are added to forward primers after both forward and reverse primer sequences are modified to support rhPCR amplification. All rhPCR primers and crRNAs were synthesized by Integrated DNA Technologies (IDT, Newark, NJ).

### Preparation of synthetic samples

Synthetic RNA samples were transcribed from gBlock DNA fragments (IDT, Newark, NJ). T7 promoter sequences included in the gBlock fragments and forward primers with T7 overhangs were used to generate templates of different sizes if needed. For *in vitro* transcription, the HiScribe® T7 High Yield RNA Synthesis Kit from New England Biolabs (NEB E2040S, Ipswich, MA) was used with the standard RNA synthesis protocol and incubated for 4 hours. After IVT was complete, IVT products were purified using a Monarch RNA Cleanup Kit (NEB, T2030L, Ipswich, MA) and stored at -80°C.

### RNA extraction and isolation

RNA was extracted from tissue culture cells using either the EZ-10 spin column kit (Bio Basic, Markham, Canada) or the MagMAX mirVana Total RNA Isolation Kit (Thermo Fisher Scientific A27828, Waltham, MA). For EZ-10 column extractions, cells were detached at their respective timepoints and lysed in 450 μL of Buffer RLT after removal of supernatant. Cell lysates were then processed following the instructions in the EZ-10 manual. For timecourse experiments involving more than 48 samples, the MagMAX mirVana kit was used in conjunction with the KingFisher extraction platform. After each timepoint, cells were immediately lysed in MagMAX Lysis/Binding Solution Concentrate (Thermo Fisher Scientific AM8500, Waltham, MA) and left in storage at -20°C until all timepoints were collected and ready for automated extraction. RNA was eluted in 50 μL of water and further processed with TURBO DNase (Thermo Fisher AM2238, Waltham, MA). RNA samples were then purified using RNAClean XP (Beckman Coulter A63987, Brea, CA) SPRI beads and eluted in 22 μL of water. Purified RNA extracts were then aliquoted and stored at -80°C until ready for usage.

### Interferon stimulation experiments

Naïve U2OS cells were cultured in 24-well cell culture plates and grown to 80% confluency prior to interferon stimulation. Cells were then stimulated with 120 IU/mL of IFN-β (Sigma-Aldrich IF014, St. Louis, MO), 800 IU/mL of IFN-α (Sigma-Aldrich SRP4594, St. Louis, MO), and 40 ng/mL of IFN-λ (R&D Systems 5259-IL, Minneapolis, MN). Stimulated cells were then incubated at 37°C for 72 hours and timepoint samples were collected for IFN-stimulated and mock wells at 0, 1, 2, 4, 8, 24, 48, and 72-hour timepoints. At each timepoint, cells were immediately lysed in MagMAX Lysis/Binding Solution Concentrate (Thermo Fisher Scientific AM8500, Waltham, MA) and stored at -20°C until all timepoint samples were ready for RNA extraction. Extractions were performed using the MagMAX mirVana kit (Thermo Fisher Scientific, Waltham, MA) and automated using the KingFisher extraction system. Extracts were treated with TURBO-DNase and further purified using RNAClean XP (SPRI) beads. Final RNA extracts were then aliquoted and stored at -80°C until required.

### Generation of virus stocks

Note that virus stocks can be obtained from Princeton University under a Uniform Biological Material Transfer Agreement (UBMTA).

#### Hepatitis C and YFV-17D

HCV RNA and subsequent viral stocks were produced as previously described^31^. In brief, viral RNA was produced via *in vitro* transcription of an XbaI-linearized HCV-Jc1 or AflII-linearized YFV-17D plasmid using the HiScribe® T7 High Yield RNA Synthesis Kit (HCV, New England Biolabs E2040S) or mMESSAGE mMACHINE™ SP6 Transcription Kit (YFV-17D, Invitrogen, AM1340) as outlined in the user manual^32^. Viral RNA was purified using the MEGAclear™ Transcription Clean-Up Kit (Invitrogen AM1908) following manufacturer’s instructions, and quality control was performed by gel electrophoresis to ensure no significant RNA degradation. Viral RNA stocks were stored as 5μg aliquots at −80°C. RNA was electroporated into Huh7.5-1(HCV) or Huh7.5 (YFV-17D) cells. Pelleted cells were resuspended in the appropriate volume of cold DPBS to achieve a concentration of 1.5e7 cells/mL. 6e6 cells were then electroporated in a 2 mm path length electroporation cuvette (BTX Harvard Apparatus; Holliston, MA) with 5μg (HCV) or 1.5μg (YFV-17D) of viral RNA using an ECM 830 Square Wave Electroporation System (BTX) at the following settings: five pulses, 99μs per pulse, 1.1 second pulse intervals, 860V. Following a ten-minute incubation at room temperature, the electroporated cells were seeded into p150 cell culture dishes and maintained in 5% (v/v) FBS DMEM. Media was changed one day post-electroporation. For HCV supernatants were collected twice daily from days four through six and stored at 4°C. For YFV-17D, supernatants were collected daily since cytopathic effects were observed to complete cell death. The pooled supernatants were passed through a 0.22μm stir cell filter (HCV) or 0.45μm syringe filter (YFV-17D) and subsequently concentrated to ∼40 mL in 100kDa MWCO Amicon® Ultra-15 Centrifugal Filter Units (Millipore Sigma UFC9100). For HCV, the TCID50/mL^33^ of concentrated virus was determined after one freeze-thaw by limiting dilution assay. For YFV-17D, virus titers were determined by focus forming assay using flavivirus group antigen antibody anti-E (clone 4G2, diluted 1:1000, 4G2, Novus Biologicals).

#### Usutu, Dengue, Zika, Langat

Infectious ZIKV (strain MR766) was kindly provided by Matthew Evans (Mount Sinai School of Medicine). Infectious LGTV was kindly provided by Marshall Bloom (NIAID). USUV (TC508) was obtained from Kenneth Plante at the World Reference Center for Emerging Viruses and Arboviruses (WRCEVA, UTMB, TX). Viral infectious cDNAs of DENV1 (FSE 2133), 2 (DAK HD 76395), 3 (FPA 0099), and 4 (BE H 403714) were generated via circular polymerase extension reaction (CPER) as previously described^34^. 7 DNA fragments covering the full-length genome and a linker fragment (containing CMV promoter, hepatitis delta virus ribozyme, and late SV40 pA signal) were generated using Q5 High-Fidelity DNA Polymerase (NEB, Ipswich, MA) from sequence-confirmed TOPO plasmids. To generate circular infectious cDNAs, equimolar quantities of each fragment were mixed to allow the overlapping region to prime the polymerase extension reaction using PrimeSTAR® GXL DNA Polymerase (TaKaRa R050A, Kusatsu, Shiga, Japan). These CPER products were then transfected into Huh7.5 cells using X-tremeGENE™ HP DNA (Roche 6366236001, Basel, Switzerland).

### Flavivirus infections

24 hours prior to infection, Huh7 cells were seeded in a 6-well format at densities of 1.5e5 (for HCV) or 6e5 cells (for all other viruses) per well. Infections were conducted in triplicate wells at an MOI of 2 (HCV), 1 (HCV), 0.1 (DENVs, ZIKV, LGTV, USUV) or 0.05 (YFV-17D). Viral inoculum was applied for 8 hours (HCV) or 2 hours (flaviviruses) after which the wells were washed once with unsupplemented DMEM and the media changed to 10% (v/v) FBS 1% (v/v) P/S DMEM. Cells were trypsinized at 2 days post-infection (dpi) (YFV-17D), 3 dpi (DENV-1, DENV-4, ZIKV, USUV, LGTV) or 5 dpi (HCV, DENV2, DENV3) and split into two tubes for either viral antigen staining or RNA extractions using the EZ-10 Spin Column Total RNA Miniprep Super Kit (Bio Basic).

### Quantification of infection

Cells were pelleted, fixed with 4% paraformaldehyde (PFA) (Sigma-Aldrich) and permeabilized in 0.1% (w/v) Saponin and 1% (v/v) FBS in DPBS. Pellets were subsequently incubated for one hour at room temperature with flavivirus group antigen antibody (clone 4G2, diluted 1:100, Novus Biologicals) for flavivirus-infected cells and murine-produced Clone 9E10 primary antibody specific for NS5A (kindly provided by Charles Rice at The Rockefeller University) diluted 1:8000 in FACS buffer (1% FBS (v/v) in DPBS) for HCV^31,35^. Cells were then washed with FACS buffer and incubated at 4°C for 30 minutes in the dark with secondary antibody (diluted 1:250, Invitrogen A-21235, Waltham, MA) conjugated with FITC (flaviviruses) or Alexa 647 (HCV). Cells were subsequently pelleted, washed once with FACS buffer and then analyzed in FACS buffer on a BD LSRII flow cytometer (BD Biosciences, Franklin Lakes, N). Flow cytometry data were processed in FlowJo Software version 10.4.2 (FlowJo, Becton Dickson, Franklin Lake, NJ).

### Epithelial-mesenchymal transition induction

MCF7 and T47D cell lines were perturbed using 5 ng/mL of recombinant human TGF-β (R&D Systems 240-B-002/CF, Minneapolis, MN) over the course of three days. 12 hours prior to treatment with TGF-β, cells were serum-starved using DMEM supplemented with 1% (v/v) FBS and 1% (v/v) P/S to improve uptake of TGF-β. After serum starvation, cells were treated with 5 ng/mL of recombinant TGF-β over 72 hours and timepoints were collected at 0, 1, 2, 4, 8, 24, 48, and 72 hours. RNA extractions were performed using the same MagMax protocol as described in the IFN stimulation experiment section above.

### cDNA synthesis

After RNA extraction and isolation, samples were converted to cDNA using SuperScript IV (Thermo Fisher Scientific 18090010, Waltham, MA) in 20 μL reactions. First, 5 μL of template RNA at 20-200 ng/μL were added to 6 μL of water, 1 μL of 50 mM oligo(dT)_20_ (Thermo Fisher Scientific 18418020, Waltham, MA), and 1 μL of 10 mM dNTP mix (NEB N0447S, Ipswich, MA). This initial premix was heated at 65°C for 5 minutes, then incubated on ice for 1 minute. A second pre-mix consisting of 4 μL of 5X SSIV buffer, 1 μL of 100 mM DTT, 1 μL of murine RNase inhibitor (NEB M0314S), 0.5 μL of SuperScript IV (Thermo Fisher Scientific 18090010, Waltham, MA), and 0.5 μL of water was prepared. After incubation on ice of the first master mix was complete, the second pre-mix (7 μL in total) was added to the 13 μL of the first mix, creating a 20 μL reaction volume. The final reaction mixture was incubated at 55°C for 20 minutes and inactivated at 80°C for 10 minutes.

### RNase H2-dependent PCR amplification

RNA samples were pre-amplified prior to Cas13 detection experiments by adding 2 μL of input material to a 25 μL reaction consisting of AmpliTaq DNA polymerase (Thermo Fisher Scientific N8080152, Waltham, MA), RNase H2 enzyme (IDT 11-03-02-02, Newark, NJ), and GEN1 rhPCR primers. GEN1 rhPCR primers consist of the conventional primer sequence with a terminal blocking moiety that takes the form of rDDDDMx, where D represents a DNA base, r represents the RNA base, M represents a mismatched DNA base, and x represents the blocker.

### Assay and sample mix preparation

Assay mixes were prepared in 16 μL volumes for each unique crRNA target consisting of 1 μL of 50 U/μL NxGen T7 Polymerase (Biosearch Technologies 30223-1), 2 μL of 800 nM LwaCas13a (Genscript), 1 μL of a 1.6 μM target-specific Cas13 crRNA, 8 μL of 2X Fluidigm Assay Loading Buffer, and 4 μL of nuclease-free water. 1.5X sample premixes were created with 2.4 μL of 10X T7 Buffer (Biosearch Technologies F88905-1, Hoddesdon, United Kingdom), 3.2 μL of 7.5X sample buffer, 0.2 μL of murine RNase inhibitor (NEB M0314S, Ipswich, MA), 0.96 μL of 25 mM rNTP mix (NEB N0466S, Ipswich, MA), 0.12 μL of 100 μM 6U FAM reporter (IDT), 0.48 μL of 50X ROX reference dye (Thermo Fisher Scientific 12223012, Waltham, MA), 1.20 μL of 20X GE buffer, and 7.44 μL of nuclease-free water. 7.5X Sample Buffer solutions were made in 5 mL aliquots with 375 μL of Tris-HCl, 75 μL of 5M NaCl, 37.5 μL of 1M MgCl_2_, 750 mg of PEG-8000, and 4512.5 μL of nuclease-free water. Sample premixes were added to pre-amplified sample mixtures at a 2:1 ratio of premix to sample.

### Integrated Fluidic Circuit (IFC) chip loading

192.24 IFCs (Standard BioTools 100-6266, South San Francisco, CA) were used for microfluidic mixing of assay and sample mixes. 3.5 μL of each sample mix were loaded into each IFC sample well and 3.5 μL of each assay mix were loaded into IFC assay wells. Remaining empty IFC wells were filled with either an assay or sample filler mix. Assay filler mix consisted of assay loading buffer and water in a 1:1 ratio and sample filler mixes consisted of nuclease-free water, 20X GE buffer, and ROX reference dye in a 93:5:2 ratio. Actuation and pressure fluid (Standard BioTools 100-6267, South San Francisco, CA) were added to the chip. 192.24 IFCs were then primed and loaded in the Fluidigm HX Controller using the default prime and load script for 30 minutes. After priming, IFCs were loaded immediately into the Biomark HD (Standard BioTools, South San Francisco, CA) to initiate the reaction. Reactions were run for three hours at 37°C with 37 imaging steps on the fluorescein amidite (FAM) channel separated by 5-minute intervals.

### Model fitting and data analysis

Parameters were generated for the 11-parameter Cas13 reaction model by fitting the model on fluorescence kinetics data from dilutions of RNA extracted from YFV17D-infected and IFN-stimulated cells. Sets of random parameters were generated and used to fit the model on dilution curves for multiple genes. A “shared fitting” approach was used to find fixed model parameters that were shared between dilutions of a given RNA target with only the input concentration parameter across dilutions. Parameter sets that successfully fit real-world fluorescence data were collected and averaged across dilutions and targets to create an initial parameter set for future model fits and analyses.

To determine input concentrations from experimental data, the initial parameter set was used to generate fits to fluorescence data using the qCARMEN model. Input concentrations were determined for targets of interest as well as two housekeeping genes, GAPDH and HPRT1. Input concentrations for all samples were normalized to housekeeping gene concentrations. Normalized concentrations were used to determine kinetic fold-changes in time-course experiments by determining relative changes from the initial timepoint. For infection data, where multiple timepoints were not collected, normalized concentrations for treated samples were compared to normalized concentrations for mock samples to determine changes in target expression.

## Data Availability

All primary data are available via Princeton DataSpace: https://doi.org/10.34770/7kbf-rv39.

## Code Availability

Source code for custom assay design and data analysis code can be found on GitHub: https://github.com/Myhrvold-Lab/qCARMEN. Automated design and analysis pipelines can be found on the LatchBio platform (link to workflows on GitHub).

## Acknowledgements

Huh7.5 cells, the anti-HCV NS5A (clone 9E10) and the pACNR-FLYF-17D-RL plasmid were kindly provided by Charles Rice (The Rockefeller University). Infectious Zika virus (MR766) was kindly provided by Matthew Evans (Mount Sinai School of Medicine). Marshall Bloom (NIAID) kindly provided the infectious Langat virus. Usutu virus (TC508) clinical isolate was kindly provided by Kenneth Plante, Ph.D. (World Reference Center for Emerging Viruses and Arboviruses at the University of Texas Medical Branch). We thank Christina DeCoste and Gabriel Palmieri at the Molecular Biology Flow Cytometry Resource Facility for excellent technical support. We thank Yibin Kang (Princeton University) for expert advice on the EMT experiments and for providing the MCF7 and T47D breast cancer cell lines.

## Author contributions

B.K., A.P. and C.M. conceived the idea. B.K. performed all qCARMEN experiments; generated all interferon stimulation and EMT samples; analyzed the data; and wrote the assay design, reaction modeling, and data analysis code. A.P. and C.M. analyzed data. J.Z., M.P.S., A.N.N., E.S. performed viral infections. J.Z. and A.G. performed qPCR experiments for Fig. 1. B.K., A.P. and C.M. wrote the paper with input from all authors.

## Extended Data Figures

**Extended Data Fig. 1.**
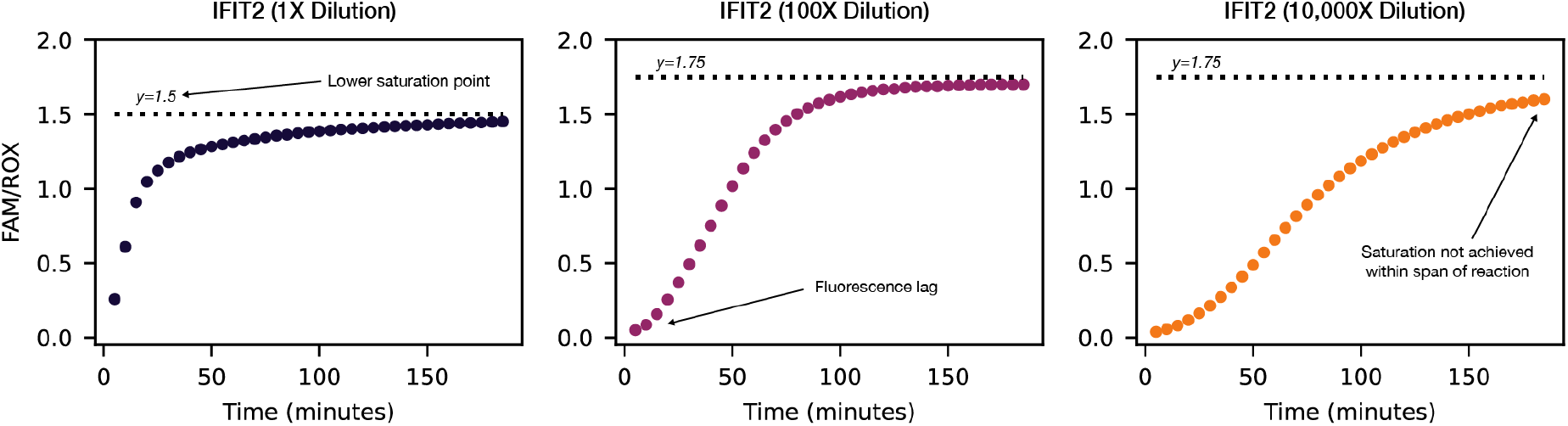
Diversity of fluorescence kinetics. From left to right: 100X serial dilutions of Huh7.5 RNA extract.

**Extended Data Fig. 2.**
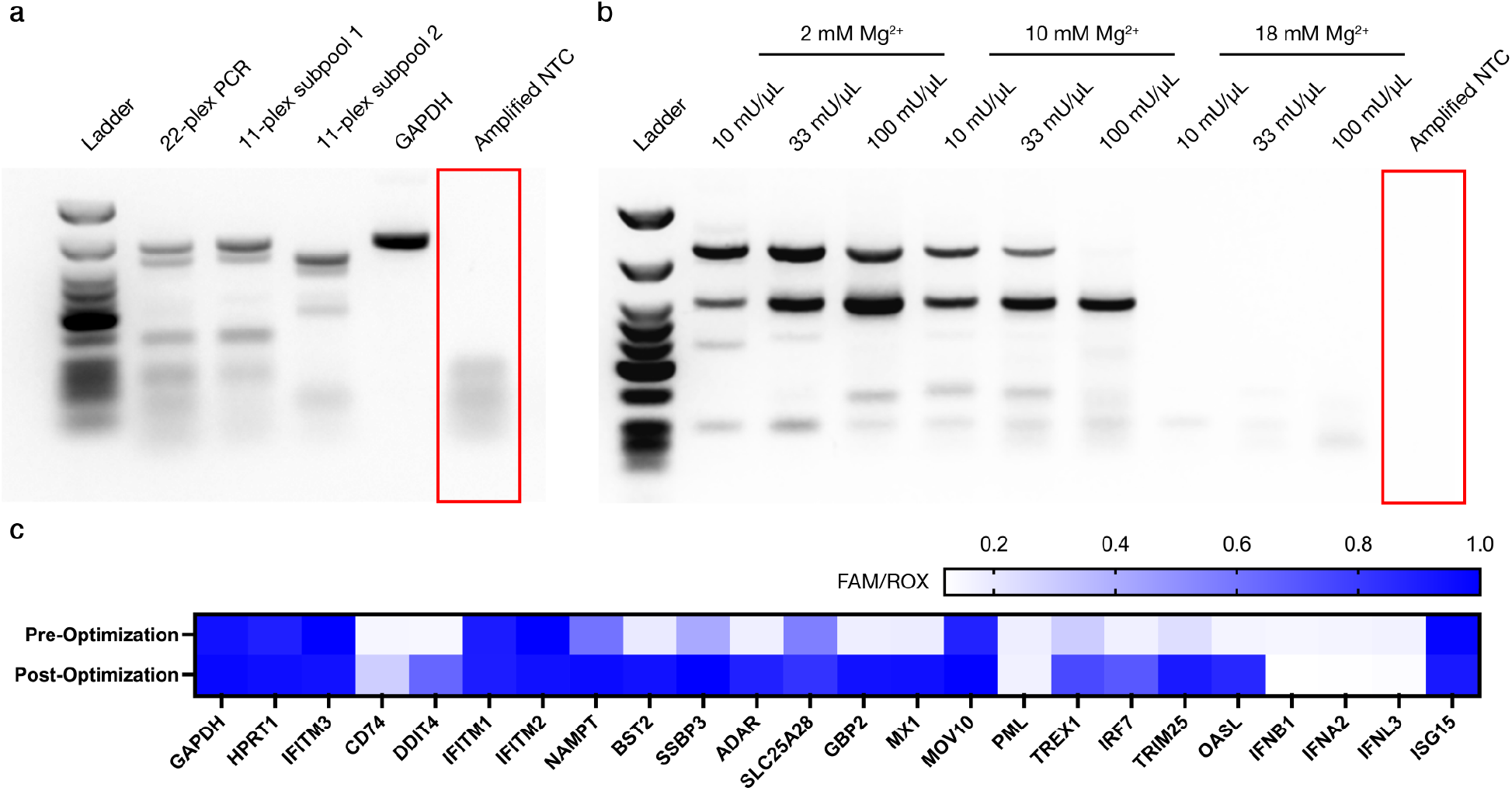
Improving multiplexed amplification with rhPCR. **a**, Multiplexed amplification of RNA extract from naive Huh7 cells with conventional PCR primers. **b**, Multiplexed amplification of RNA extract with rhPCR primers. **c**, Fluorescence output after 3 hours of qCARMEN reaction for ISG panel.

**Extended Data Fig. 3.**
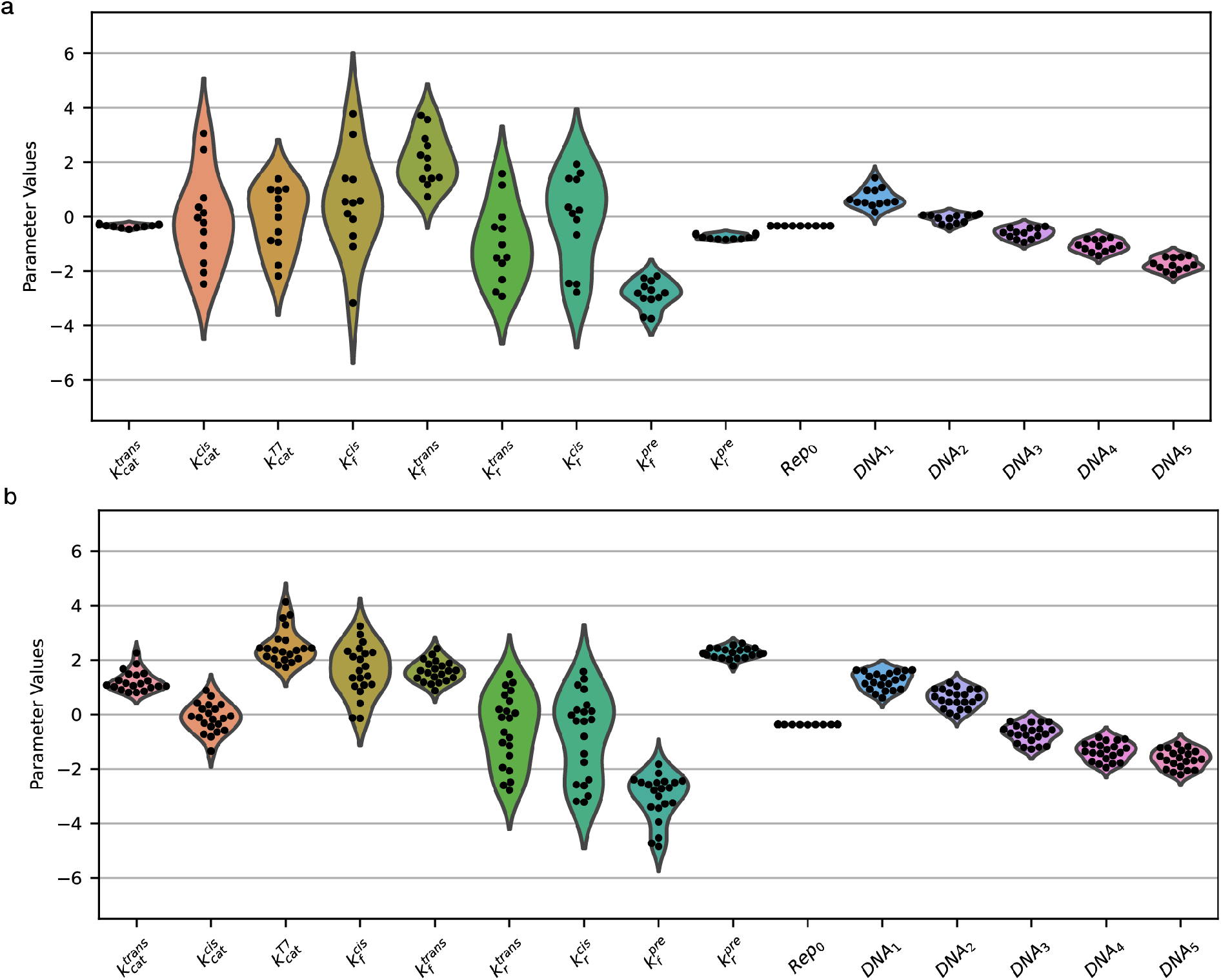
Model parameter distributions. **a**, Calculated parameter values for GAPDH. **b**, Calculated parameter values for ADAR.

**Extended Data Fig. 4.**
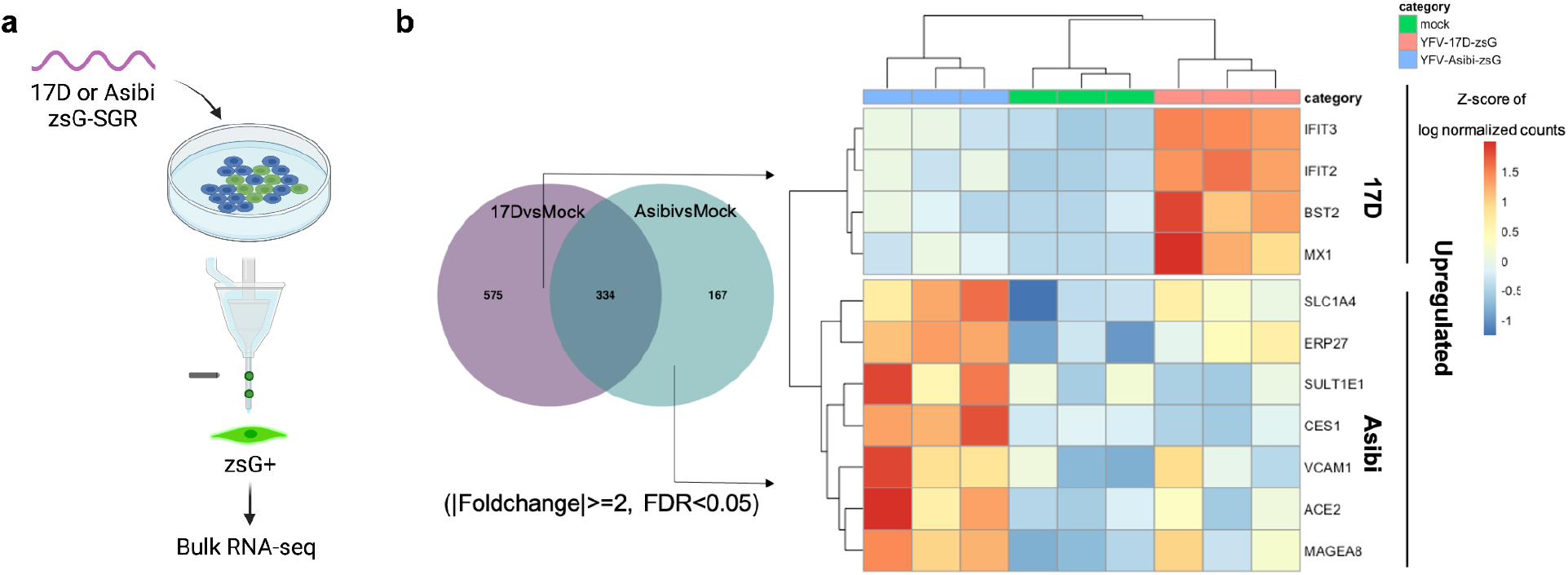
RNA-Seq analysis of Huh7 cells harboring subgenomic YFV17D and YFV-Asibi replicons. **a**, RNA-Seq workflow. **b**, Summary of uniquely upregulated innate immunity genes.

**Extended Data Fig. 5.**
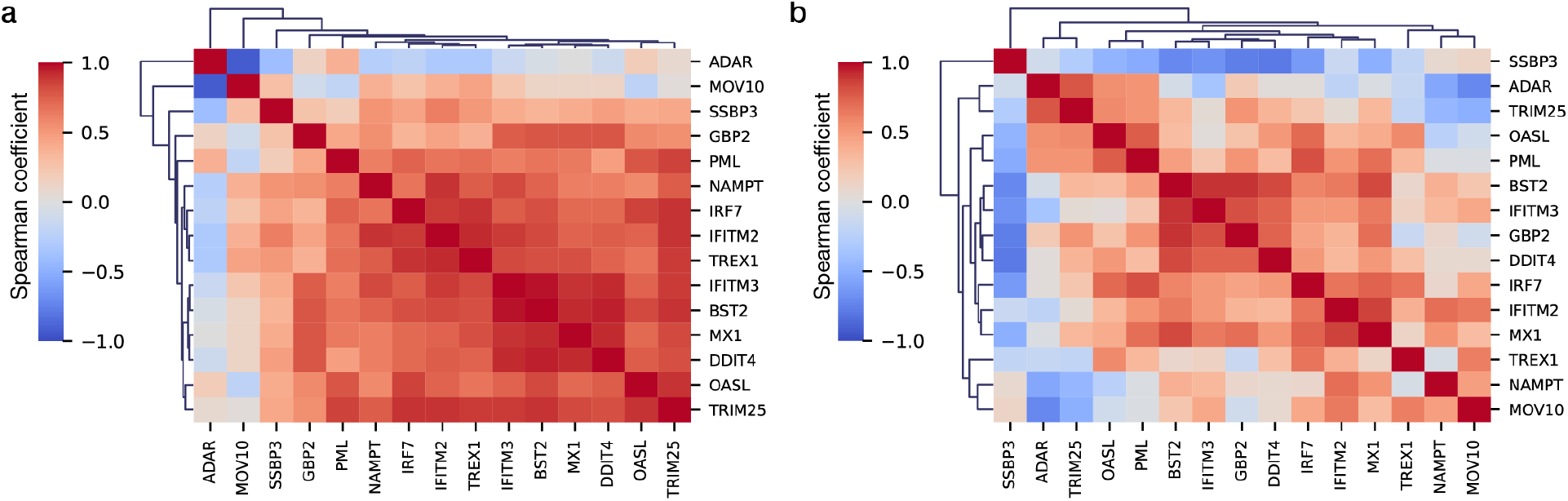
Spearman correlations with interferon response kinetics. **a**, Spearman correlations for IFN-α stimulation experiments. **b**, Spearman correlations for IFN-λ stimulation results.

**Extended Data Fig. 6.**
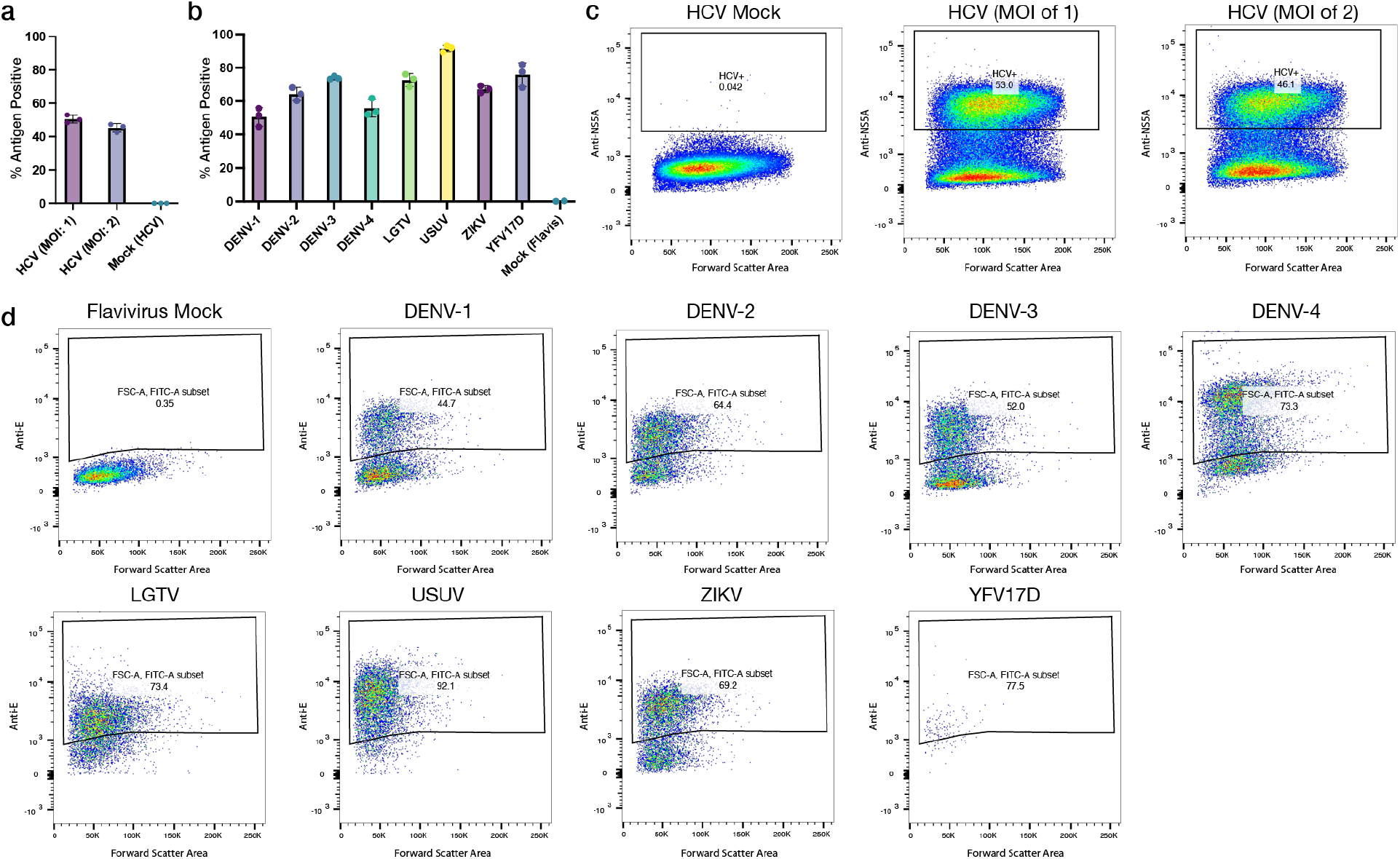
Antigen staining for infection quantification. **a**, Percent of antigen-positive cells for HCV (2 MOIs). **b**, Percent of antigen-positive cells for flavivirus-infected cells. Error bars presented as SD. **c**, Infected cell populations for HCV samples. **d**, Infected cell populations for flavivirus samples.

