## Supplementary Information for "Highly multiplexed mRNA quantitation with CRISPR-Cas13"

PDF contains:

Supplementary Tables 1-7
Supplementary Notes 1-2

| | gBlock Sequence |
|---|---|
| GAPDH | GCTCTCTGCTCCTCCTGTTCGACAGTCAGCCGCATCTTCTTTTGCGTCGC<br>CAGCCGAGCCACATCGCTCAGACACCATGGGGAAGGTGAAGGTCGGAG<br>TCAACGGATTTGGTCGTATTGGGCGCCTGGTCACCAGGGCTGCTTTTAA<br>CTCTGGTAAAGTGGATATTGTTGCCATCAATGACCCCTTCATTGACCTC<br>AACTACATGGTTTACATGTTCCAATATGATTCCACCCATGGCAAATTCC<br>ATGGCACCGTCAAGGCTGAGAACGGGAAGCTTGTCATCAATGGAAATC<br>CCATCACCATCTTCCAGGAGCGAGATCCCTCCAAAATCAAGTGGGGCG<br>ATGCTGGCGCTGAGTACGTCGTGGAGTCCACTGGCGTCTTCACCACCAT<br>GGAGAAGGCTGGGGCTCATTTGCAGGGGGGAGCCAAAAGGGTCATCAT<br>CTCTGCCCCCTCTGCTGATGCCCCCATGTTCGTCATGGGTGTGAACCAT<br>GAGAAGTATGACAACAGCCTCAAGATCATCAGCAATGCCTCCTGCACC<br>ACCAACTGCTTAGCACCCCTGGCCAAGGTCATCCATGACAACTTTGGTA<br>TCGTGGAAGGACTCATGACCACAGTCCATGCCATCACTGCCACCCAGA<br>AGACTGTGGATGGCCCCTCCGGGAAACTGTGGCGTGATGGCCGCGGGG<br>CTCTCCAGAACATCATCCCTGCCTCTACTGGCGCTGCCAAGGCTGTGGG<br>CAAGGTCATCCCTGAGCTGAACGGGAAGCTCACTGGCATGGCCTTCCGT<br>GTCCCCACTGCCAACGTGTCAGTGGTGGACCTGACCTGCCGTCTAGAAA<br>AACCTGCCAAATATGATGACATCAAGAAGGTGGTGAAGCAGGCGTCGG<br>AGGGCCCCCTCAAGGGCATCCTGGGCTACA |
| HPRT1 | ATTTTATCAGACTGAAGAGCTATTGTAATGACCAGTCAACAGGGGACAT<br>AAAAGTAATTGGTGGAGATGATCTCTCAACTTTAACTGGAAAGAATGTC<br>TTGATTGTGGAAGATATAATTGACACTGGCAAAACAATGCAGACTTTGC<br>TTTCCTTGGTCAGGCAGTATAATCCAAAGATGGTCAAGGTCGCAAGCTT<br>GCTGGTGAAAAGGACCCCACGAAGTGTTGGATATAAGCCAGACTTTGTT<br>GGATTTGAAATTCCAGACAAGTTTGTTGTAGGATATGCCCTTGACTATA<br>ATGAATACTTCAGGGATTTGAATCATGTTTGTGTCATTAGTGAAACTGG<br>AAAAGCAAAATACAAAGCCTAAGATGAGAGTTCAAGTTGAGTTTGGAA<br>ACATCTGGAGTCCTATTGACATCGCCAGTAAAATTATCAATGTTCTAGT<br>TCTGTGGCCATCTGCTTAGTAGAGCTTTTTGCATGTATCTTCTAAGAATT<br>TTATCTGTTTTGTACTTTAGAAATGTCAGTTGCTGCATTCCTAAACTGTT<br>TATTTGCACTATGAGCCTATAGACTATCAGTTCCCTTTGGGCGGATTGTT<br>GTTTAACTTGTAAATGAAAAAATTCTCTTAAACCACAGCACTATTGAGT<br>GAAACATTGAACTCATATCTGTAAGAAATAAAGAGAAGATATATTAGT<br>TTTTTAATTGGTATTTTAATTTTTATATATGCAGGAAAGAATAGAAGTG<br>ATTGAATATTGTTAATTATACCACCGTGTGTTAGAAAAGTAAGAAGCAG<br>TCAATTTTCACATCAAAGACAGCATCTAAGAAGTTTTGTTCTGTCCTGG<br>AATTATTTTAGTAGTGTTTCAGTAATGTTGACTGTATTTTCCAACTTGTT<br>CAAATTATTACCAGTGAATCTTTGTCAGCAGTTCCCTTTTAAATGCAAAT<br>CAATAAATTCCCAAAAATTTAA |
| FGFR2 IIIb | CCACGGTCCGCTACCTTACAGGAATTGAGACCGTCCATTAATTTCCCTT<br>GCATTTTCATCGGGCCGTGCAGGCTAATACGACTCACTATAGGGAAATA<br>agccaccaaccaaataccaaatctctcaaccagaagtgtacgtggctgcgccaggggagtcgctagaggtgcgctgc<br>ctgttgaaagatgccgccgtgatcagttggactaaggatggggtgcacttggggcccaacaataggacagtgcttattg<br>gggagtacttgcagataaagggcgccacgcctagagactccggcctctatgcttgtactgccagtaggactgtagaca |

| | |
|---|---|
| | gtgaaacttggtacttcatggtgaatgtcacagatgccatctcatccggagatgatgaggatgacaccgatggtgcgga<br>agattttgtcagtgagaacagtaacaacaagagagcaccatactggaccaacacagaaaagatggaaaagcggctcc<br>atgctgtgcctgcggccaacactgtcaagtttcgctgcccagccgggggggaacccaatgccaaccatgcggtggctg<br>aaaaacgggaaggagtttaagcaggagcatcgcattggaggctacaaggtacgaaaccagcactggagcctcattat<br>ggaaagtgtggtcccatctgacaagggaaattatacctgtgtagtggagaatgaatacgggtccatcaatcacacgtac<br>cacctggatgttgtggagcgatcgcctcaccggcccatcctccaagccggactgccggcaaatgcctccacagtggtc<br>ggaggagacgtagagtttgtctgcaaggtttacagtgatgcccagccccacatccagtggatcaagcacgtggaaaag<br>aacggcagtaaatacgggcccgacgggctgccctacctcaaggttctcaagcactcggggataaatagttccaatgca<br>gaagtgctggctctgttcaatgtgaccgaggcggatgctggggaatatatatgtaaggtctccaattatatagggcaggc<br>caaccagtctgcctggctcactgtcctgccaaaacagcaagcgcctggaagagaaaaggagattacagcttccccag<br>actacctggagatagccatttactgcataggggtcttcttaatcgcctgtatggtggtaacagtcatcctgtgccgaatgaa<br>gaacacgaccaagaagccagacttcagcagccagccggctgtgcacaagctgaccaaacgtatccccctgcggaga<br>caggtaacagtttcggctgagtccagctcctccatgaactccaacaccccgctggtgaggataacaacacgcctctctt<br>caacggcagacacccccatgctggcaggggtctccgagtatgaacttccagaggacccaaaatgggagtttccaaga<br>gataagctgacactgggcaagcccctgggagaaggttgctttgggcaagtggtcatggcggaagcagtgggaattga<br>caaagacaagcccaaggaggcggtcaccgtggccgtgaagatgttgaaagatgatgccacagagaaagacctttctg<br>atctggtgtcagagatggagatgatgaagatgattgggaaacacaagaatatcataaatcttcttggagcctgcacacag<br>gatgggcctctctatgtcatagttgagtatgcctctaaaggcaacctccgagaatacctccgagcccggaggccacccg<br>ggatggagtactcctatgacattaaccgtgttcctgaggagcagatgaccttcaaggacttggtgtcatgcacctaccag<br>ctggccagaggcatggagtacttggcttcccaaaaa |
| FGFR2<br>IIIc | CCACGGTCCGCTACCTTACAGGAATTGAGACCGTCCATTAATTTCCCTT<br>GCATTTTCATCGGGCCGTGCAGGCTAATACGACTCACTATAGGGAAATA<br>agccaccaaccaaataccaaatctctcaaccagaagtgtacgtggctgcgccaggggagtcgctagaggtgcgctgc<br>ctgttgaaagatgccgccgtgatcagttggactaaggatggggtgcacttggggcccaacaataggacagtgcttattg<br>gggagtacttgcagataaagggcgccacgcctagagactccggcctctatgcttgtactgccagtaggactgtagaca<br>gtgaaacttggtacttcatggtgaatgtcacagatgccatctcatccggagatgatgaggatgacaccgatggtgcgga<br>agattttgtcagtgagaacagtaacaacaagagagcaccatactggaccaacacagaaaagatggaaaagcggctcc<br>atgctgtgcctgcggccaacactgtcaagtttcgctgcccagccgggggggaacccaatgccaaccatgcggtggctg<br>aaaaacgggaaggagtttaagcaggagcatcgcattggaggctacaaggtacgaaaccagcactggagcctcattat<br>ggaaagtgtggtcccatctgacaagggaaattatacctgtgtagtggagaatgaatacgggtccatcaatcacacgtac<br>cacctggatgttgtggagcgatcgcctcaccggcccatcctccaagccggactgccggcaaatgcctccacagtggtc<br>ggaggagacgtagagtttgtctgcaaggtttacagtgatgcccagccccacatccagtggatcaagcacgtggaaaag<br>aacggcagtaaatacgggcccgacgggctgccctacctcaaggttctcaaggccgccggtgttaacaccacggacaa<br>agagattgaggttctctatattcggaatgtaacttttgaggacgctggggaatatacgtgcttggcgggtaattctattggg<br>atatcctttcactctgcatggttgacagttctgccagcgcctggaagagaaaaggagattacagcttccccagactacct<br>ggagatagccatttactgcataggggtcttcttaatcgcctgtatggtggtaacagtcatcctgtgccgaatgaagaacac<br>gaccaagaagccagacttcagcagccagccggctgtgcacaagctgaccaaacgtatccccctgcggagacaggta<br>acagtttcggctgagtccagctcctccatgaactccaacaccccgctggtgaggataacaacacgcctctcttcaacgg<br>cagacacccccatgctggcaggggtctccgagtatgaacttccagaggacccaaaatgggagtttccaagagataag<br>ctgacactgggcaagcccctgggagaaggttgctttgggcaagtggtcatggcggaagcagtgggaattgacaaaga<br>caagcccaaggaggcggtcaccgtggccgtgaagatgttgaaagatgatgccacagagaaagacctttctgatctggt<br>gtcagagatggagatgatgaagatgattgggaaacacaagaatatcataaatcttcttggagcctgcacacaggatggg<br>cctctctatgtcatagttgagtatgcctctaaaggcaacctccgagaatacctccgagcccggaggccacccgggatgg<br>agtactcctatgacattaaccgtgttcctgaggagcagatgaccttcaaggacttggtgtcatgcacctaccagctggcc<br>agaggcatggagtacttggcttcccaaaaa |

**Supplementary Table 1:** gBlock sequences for synthetic RNA targets.

| | Direction | Sequence |
|---|---|---|
| GAPDH | Forward | TAATACGACTCACTATAGGGATAATAGTTGCCATCAATGACCCCTTCATTG |
| GAPDH | Reverse | ATGACGAACATGGGGGCATCAG |
| HPRT1 | Forward | TAATACGACTCACTATAGGGATAATCTTCAGGGATTTGAATCATGTTTGTGTCAT |
| HPRT1 | Reverse | TTCACTCAATAGTGCTGTGGTTTAAGAGAA |

**Supplementary Table 2:** PCR primers for incorporation of T7 promoter overhang into gBlock targets.

| | **Forward Primer** | **Reverse Primer** |
|---|---|---|
| GAPDH | CACCATCTTCCAGGAGCGAGA | ATGACGAACATGGGGGCATCAG |
| HPRT1 | ACCAGTCAACAGGGGACATAA | CTTCGTGGGGTCCTTTTCACC |
| IFIT2 | GGGAAACTATGCCTGGGTC | CCTTCGCTCTTTCATTTTGGTTTC |
| IFIT3 | TCAGAAGTCTAGTCACTTGGGG | ACACCTTCGCCCTTTCATTTC |
| MX1 | GTTTCCGAAGTGGACATCGCA | CTGCACAGGTTGTTCTCAGC |
| ACE2 | TCCCTGCTCATTTGCTTGGTG | ATATTCTCTGTGCATCCCAGGC |
| SULT1E1 | GCCGGAATGCAAAGGATGTG | AGGAACCATAAGGAACCTGTCC |
| CES1 | ACCCCTGAGGTTTACTCCACC | TGCACATAGGAGGGTACGAGG |
| CSRP1 | TAATACGACTCACTATAGCAGAATGCCGAACTGGGGAGGAG | TCTTGCAGACCATGCACAGGAAGC |
| CSRP2 | TAATACGACTCACTATAGGAGCTGGAAAGCCCTGGCACAA | AAAAGAAATGTTCCCCCAAAATGCTGGTG |
| TGFBI | TAATACGACTCACTATAGCCTTGCCAGCTGAAGTGCTGGAC | CGTGTACTGGCCGTTACCTTCAAGC |
| TUFT1 | TAATACGACTCACTATAGGTGGCTTTTGAGGTCACACTCAGCC | GGACTTTCCGCTGCTGGCTCT |

**Supplementary Table 3:** RT-qPCR primers for gene expression quantitation.

| | crRNA Sequence | qCARMEN Forward | qCARMEN Reverse |
|---|---|---|---|
| GAPDH | GAUUUAGACUACCCCAAAAACGAAGGGGACUAAAACUCAAUGAAGGGGUCAUUGAUGGCAACAA | TAATACGACTCACTATAGGGAAATACCAGGGCTGCTTTTAACTCTGGTAAAGTGGrATATTT/iSpC3/ | GGCATTGCTGATGATCTTGAGGCTGTTGTCrATACTC/iSpC3/ |
| HPRT1 | GAUUUAGACUACCCCAAAAACGAAGGGGACUAAAACGUAAUCCAGCAGGUCAGCAAAGAAUUUA | TAATACGACTCACTATAGGGAAATAACAGGACTGAACGTCTTGCTCGAGATrGTGATT/iSpC3/ | TGCCTGACCAAGGAAAGCAAAGTCTGrCATTGC/iSpC3/ |
| IFITM3 | GAUUUAGACUACCCCAAAAACGAAGGGGACUAAAACACAGGAGAGAAGAAGGUUUGGACAGUGU | TAATACGACTCACTATAGGGAAATAGCTGGTCTTCGCTGGACACCATGAATrCACACC/iSpC3/ | AACCATCTTCCTGTCCCTAGACTTCACGGArGTAGGA/iSpC3/ |
| CD74 | GAUUUAGACUACCCCAAAAACGAAGGGGACUAAAACUGUCAGGAACGGGGAUCUGUCUAGCUUU | TAATACGACTCACTATAGGGAAATACCAGTCCCCATGTGAGAGCAGCAGArGGCGGC/iSpC3/ | ATGAGCCTAGGTCTTGGAGCCTTGTGTTrGGAGGA/iSpC3/ |
| DDIT4 | GAUUUAGACUACCCCAAAAACGAAGGGGACUAAAACGGGAGAGUUGGCGGAGCUAAACAGCCCC | TAATACGACTCACTATAGGGAAATAAGGAAGACACGGCTTACCTGGATGGrGGTGTA/iSpC3/ | TTCTTGATGACTCGGAAGCCAGTGCTCArGCGTCG/iSpC3/ |
| IFITM1 | GAUUUAGACUACCCCAAAAACGAAGGGGACUAAAACUAACAUAAUAUGGUAGACUGUCACAGAG | TAATACGACTCACTATAGGGAAATACGTGAAGTCTAGGGACAGGAAGATGGTTrGGCGAA/iSpC3/ | GAATGACACTGTAGACAGGTGTGTGGGTrATAAAA/iSpC3/ |
| IFITM2 | GAUUUAGACUACCCCAAAAACGAAGGGGACUAAAACCGCUGGGCCUGGACGACCAACACUGGGA | TAATACGACTCACTATAGGGAAATACGTGAAGTCTAGGGACAGGAAGATGGTTrGGCGAA/iSpC3/ | TGTGAGGATAAAGGGCTGATGCAGGACTrCGGCTT/iSpC3/ |
| NAMPT | GAUUUAGACUACCCCAAAAACGAAGGGGACUAAAACGUACUUAUUAAGAAUGUACUGCAACCCA | TAATACGACTCACTATAGGGAAATAGCCACCGACTCCTACAAGGTTACTCACTATrAAACAG/iSpC3/ | TGGGAATGACAAAGCCCTCAGGAACArGCTTTC/iSpC3/ |
| BST2 | GAUUUAGACUACCCCAAAAACGAAGGGGACUAAAACAGACAAGAGGGGGGACUCAUUGUCCGGA | TAATACGACTCACTATAGGGAAATAATCCCAGGAAGCTGGCACATCTTGGAArGGTCCT/iSpC3/ | AACCCATAACAACAGGCAGCACATGCrCCCCCG/iSpC3/ |

| | | | |
|---|---|---|---|
| SSBP3 | GAUUUAGACUACCCCAAAAACGAAGGGGACUAAAACUGUUGUCACUGGAAUUUGUUGAAUCUGC | TAATACGACTCACTATAGGGAAATATATGGATCCCACACGACAACAAGGCCArCCCCAG/iSpC3/ | CCAGGCGGCACTGGATTAATCATTGTGTAGrATGTTT/iSpC3/ |
| ADAR | GAUUUAGACUACCCCAAAAACGAAGGGGACUAAAACUUUGGAGACCCGGGACAAUUCAUUCCGU | TAATACGACTCACTATAGGGAAATATGTGGATGGGCCACGGAATGAATTGTrCCCGGT/iSpC3/ | TGAGGAATGCTACGACCTACCTCTCTCArCACCCC/iSpC3/ |
| SLC25A28 | GAUUUAGACUACCCCAAAAACGAAGGGGACUAAAACGUUGUGGCUGCGGCAGCUACAGCUCCUG | TAATACGACTCACTATAGGGAAATAAGCCATGAACCCTGCGGAAGTGGTrCAAGCG/iSpC3/ | GGACTCCTGGGTGTTGAGCAGTGTTrUTGCAG/iSpC3/ |
| GBP2 | GAUUUAGACUACCCCAAAAACGAAGGGGACUAAAACGCUGAGGAUGUAGGAACAAAAUUCUGCA | TAATACGACTCACTATAGGGAAATAGGACCAACTTCACTATGTGACAGAGCTGACrAGATCT/iSpC3/ | CTGGAATGCCACCTGAAAGAGTCTTGACrATTGGG/iSpC3/ |
| MX1 | GAUUUAGACUACCCCAAAAACGAAGGGGACUAAAACGCAGUGAUGUCCUGAUUAAAGGCAUUAA | TAATACGACTCACTATAGGGAAATATTGAGAACCACCCATATTTCAGGGATCTGCrUGGAGT/iSpC3/ | GGAACTCGTGTCGGAGTCTGGTAAACArGCCGAG/iSpC3/ |
| MOV10 | GAUUUAGACUACCCCAAAAACGAAGGGGACUAAAACCUUAAGGAAACCCAGAUUAAAGUCCAGA | TAATACGACTCACTATAGGGAAATAGGATGACATCAAGGACTTGAAGGTGGGTTCrAGTAGG/iSpC3/ | CAGTAAATTCTGTCCCTGTTGCAGGTCCrAGTTTT/iSpC3/ |
| PML | GAUUUAGACUACCCCAAAAACGAAGGGGACUAAAACCUGGAUCUCUGCGCUGAUGUCGCACUUG | TAATACGACTCACTATAGGGAAATATGACCAGCATCTACTGCCGAGGATGTTrCCAAGA/iSpC3/ | ATCTCCTCGTAGTCGCGCTGGTACrCGCGCT/iSpC3/ |
| TREX1 | GAUUUAGACUACCCCAAAAACGAAGGGGACUAAAACCCACACACAGGGAGAGCUUGUCUACCAC | TAATACGACTCACTATAGGGAAATACCCATGCAGACCCTCATCTTTTTCGACATGrGAGGCA/iSpC3/ | GAGCAGGTTGGCCAGGTTGTCATCAAAArCATTGG/iSpC3/ |
| IRF7 | GAUUUAGACUACCCCAAAAACGAAGGGGACUAAAACCUGAGGCUGCUGCUAUCCAGGGAAGACA | TAATACGACTCACTATAGGGAAATATCTTCTTCCAAGAGCTGGTGGAATTCCGrGGCACT/iSpC3/ | GCTGCTCCAGCTTTCTGGAGTTCTCATTAGrACTGGT/iSpC3/ |

| | | | |
|---|---|---|---|
| TRIM25 | GAUUUAGACUACCCCAAAAACGAAGGGGACUAAAACGUUGUAGUCCAGGAUGACUUUAAUGUAA | TAATACGACTCACTATAGGGAAATAAGTCCAGACCTGAGCTCCTGGAGTATTArCATTAG/iSpC3/ | ATCTTGGTGTTGAACCACTCCACGCArCCAGGG/iSpC3/ |
| OASL | GAUUUAGACUACCCCAAAAACGAAGGGGACUAAAACCGAGAGCAUCGGGGACUCUCUGCUCCAU | TAATACGACTCACTATAGGGAAATATAGTCAAGGTGGGCTCCTTCGGGAATrGGCACT/iSpC3/ | AGGATGGGCAGAAATTTCCAGGACCArCCGCAT/iSpC3/ |
| IFNB1 | GAUUUAGACUACCCCAAAAACGAAGGGGACUAAAACAGACACUUGUUGGUCAUGUUGACAACAC | TAATACGACTCACTATAGGGAAATAAGCCTTTGCTCTGGCACAACAGGTAGTArGGCGAA/iSpC3/ | ATCTCATAGATGGTCAATGCGGCGTCCTrCCTTCC/iSpC3/ |
| IFNA2 | GAUUUAGACUACCCCAAAAACGAAGGGGACUAAAACGAAUUUGUCUAGGAGGGUCUCAUCCCAA | TAATACGACTCACTATAGGGAAATAAAAGGCTGAAACCATCCCTGTCCTCCATrGAGATT/iSpC3/ | ATGATTTCTGCTCTGACAACCTCCCAGGrCACAAT/iSpC3/ |
| IFNL3 | GAUUUAGACUACCCCAAAAACGAAGGGGACUAAAACGAGCGGCACUUGCAGUCCUUCAGCAGAA | TAATACGACTCACTATAGGGAAATACTTAGAAGAGTCGCTTCTGCTGAAGGACTGrCAAGTT/iSpC3/ | AAGAGGTTGAAGGTGACAGAGGCCTCrGAGGCG/iSpC3/ |
| ISG15 | GAUUUAGACUACCCCAAAAACGAAGGGGACUAAAACGUGCUGCUGCGGCCCUUGUUAUUCCUCA | TAATACGACTCACTATAGGGAAATAAGATCACCCAGAAGATCGGCGTGCArCGCCTG/iSpC3/ | TTCCCCTCGAAGGTCAGCCAGAACArGGTCGC/iSpC3/ |

**Supplementary Table 4:** ISG panel crRNAs and rhPCR primer sets.

| Gene | Internal ID | NCBI Accession IDs | | | |
|---|---|---|---|---|---|
| WASF1 | 1 | NM_003931.3 | | | |
| WASF1 | 2 | NM_001024934.2 | | | |
| WASF1 | 3 | NM_001024935.2 | | | |
| WASF1 | 4 | NM_001024936.2 | | | |
| SCRIB | 1 | NM_182706.5 | | | |
| SCRIB | 2 | NM_015356.5 | | | |
| FGFR2 | 2c | NM_000141 | | | |
| FGFR2 | 2b | NM_022970 | | | |
| FGFR1 | Beta | NM_023105 | | | |
| FGFR1 | Alpha | NM_023110 | | | |
| PPFIBP1 | 1 | NM_003622.4 | | | |
| PPFIBP1 | 2 | NM_177444.3 | | | |
| PPFIBP1 | 3 | NM_001198915.2 | | | |
| PPFIBP1 | 4 | NM_001198916.2 | | | |
| SMAD2 | 1 | NM_005901.6 | | | |
| SMAD2 | 2 | NM_001003652.4 | NM_001135937.3 | | |
| DKK3 | 1A | NM_013253.5 | NM_001330220.3 | | |
| DKK3 | 1B | NM_015881.6 | | | |
| CD44 | Standard | NM_001001391.2 | NM_001202557.2 | | |
| CD44 | Variant | NM_001001389.2 | NM_000610.4 | | |
| EXOC7 | Epithelial | NM_001375976.1 | NM_001145299.4 | NM_001145298.4 | NM_001145297.4 |
| EXOC7 | Mesenchymal | NM_001375975.1 | NM_015219.5 | | |
| FLNB | Hinge 1 | NM_001164317.2 | NM_001457.4 | | |
| FLNB | No Hinge 1 | NM_001164318.2 | NM_001164319.2 | | |
| ENAH | 11a | NM_001377481.1 | NM_001008493.3 | | |
| ENAH | No 11a exon | NM_018212.6 | NM_001377482.1 | NM_001377483.1 | |

**Supplementary Table 5:** Isoform panel IDs and corresponding accession numbers.

| Gene | Internal ID | crRNA Sequence |
| --- | --- | --- |
| WASF1 | 1 | GAUUUAGACUACCCCAAAAACGAAGGGGACUAAAACU UUAACAUGCUGAUUUCUCCAACACUAG |
| WASF1 | 2 | GAUUUAGACUACCCCAAAAACGAAGGGGACUAAAACC AGCAUAUUUACUUAGGCUACUUAGUUG |
| WASF1 | 3 | GAUUUAGACUACCCCAAAAACGAAGGGGACUAAAACC UACUUAGUUGUCUAAUUAUAUUUGCCA |
| WASF1 | 4 | GAUUUAGACUACCCCAAAAACGAAGGGGACUAAAACU UAACCUUUGUGCCAGUUCACCGUGAAA |
| SCRIB | 1 | GAUUUAGACUACCCCAAAAACGAAGGGGACUAAAACG UAGUCAAACUUCUUUCCAGACAAGGGC |
| SCRIB | 2 | GAUUUAGACUACCCCAAAAACGAAGGGGACUAAAACC GGGCCAGGGCUGGGAGAUGGUUCCAGG |
| FGFR2 | 2c | GAUUUAGACUACCCCAAAAACGAAGGGGACUAAAACU UGGUCCAGUAUGGUGCUCUCUUGUUGU |
| FGFR2 | 2b | GAUUUAGACUACCCCAAAAACGAAGGGGACUAAAACG CUGGACUCAGCCGAAACUGUUACCUGU |
| FGFR1 | Beta | GAUUUAGACUACCCCAAAAACGAAGGGGACUAAAACA GAGCAUCUUGUUCAGGCAAGGUCGGGG |
| FGFR1 | Alpha | GAUUUAGACUACCCCAAAAACGAAGGGGACUAAAACA CGGGCAUACGGUUUGGUUUGGUGUUAU |
| PPFIBP1 | 1 | GAUUUAGACUACCCCAAAAACGAAGGGGACUAAAACC UCUUCAUAUUCUCCAUCUUUGCCUUUU |
| PPFIBP1 | 2 | GAUUUAGACUACCCCAAAAACGAAGGGGACUAAAACC ACCUUCUCUCUGUGCUCUUCAAGACAA |
| PPFIBP1 | 3 | GAUUUAGACUACCCCAAAAACGAAGGGGACUAAAACG GUGUCCAUUUGUAUUUUUAAAAAGCAG |
| PPFIBP1 | 4 | GAUUUAGACUACCCCAAAAACGAAGGGGACUAAAACC UUUUAAAGAUGCCAGUUCAUCUUUGGU |
| SMAD2 | 1 | GAUUUAGACUACCCCAAAAACGAAGGGGACUAAAACA GGAAAACAGCCUCUUGUAUCGAACCUG |
| SMAD2 | 2 | GAUUUAGACUACCCCAAAAACGAAGGGGACUAAAACG UAAGGGUUUACACAUACUUCAUCCUUU |
| DKK3 | 1A | GAUUUAGACUACCCCAAAAACGAAGGGGACUAAAACG UUGCGCUGCGGGCCGGACUGGAUCUGC |
| DKK3 | 1B | GAUUUAGACUACCCCAAAAACGAAGGGGACUAAAACG ACCAAGCACAGGUCAGCCCCCUCCCCU |
| CD44 | Standard | GAUUUAGACUACCCCAAAAACGAAGGGGACUAAAACC UCUGGUAGCAGGGAUUCUGUCUGUGCU |
| CD44 | Variant | GAUUUAGACUACCCCAAAAACGAAGGGGACUAAAACC CAGAAAAACUGAGGUGUCUGUCUCUUU |
| EXOC7 | Epithelial | GAUUUAGACUACCCCAAAAACGAAGGGGACUAAAACC ACUGACGCAGUGGAUGUAGGCAUCGGU |
| EXOC7 | Mesenchymal | GAUUUAGACUACCCCAAAAACGAAGGGGACUAAAACG UAGACGUUCAUGAAAUCUUGGUUGCGG |

| | | |
|---|---|---|
| FLNB | Hinge 1 | GAUUUAGACUACCCCAAAAACGAAGGGGACUAAAACCAAACGGAAUGACCAGGUCAAAAGGCUU |
| FLNB | No Hinge 1 | GAUUUAGACUACCCCAAAAACGAAGGGGACUAAAACUCACACUUAAUGCCAACACUGACAUCAC |

**Supplementary Table 6:** Isoform panel crRNA sequences.

| Gene | Isoform ID | Direction | Sequence |
|---|---|---|---|
| WASF1 | 1 | fw | TAATACGACTCACTATAGGGAAATAAGCTTCGGTACTCTGACACCTTCTCTTGrCACTTT/iSpC3/ |
| WASF1 | 1 | rev | GGCACAAGTGCCTAGGATCGATGTTTCTrUTTCAA/iSpC3/ |
| WASF1 | 2 | fw | TAATACGACTCACTATAGGGAAATACCAGCTAGACTAGGTGAACTGGCACAAArGGTTAG/iSpC3/ |
| WASF1 | 2 | rev | GGTCCACACGTTCTTGCAATGAGTTGACrUCTGAG/iSpC3/ |
| WASF1 | 3 | fw | TAATACGACTCACTATAGGGAAATAAGCCCCCAAGATTTCACGTGTTGGAGAArATCAGA/iSpC3/ |
| WASF1 | 3 | rev | GGTCCACACGTTCTTGCAATGAGTTGACrUCTGAG/iSpC3/ |
| WASF1 | 4 | fw | TAATACGACTCACTATAGGGAAATAGACCGCTGCCGAAGCATGAAGAArGGGGTG/iSpC3/ |
| WASF1 | 4 | rev | GGTCCACACGTTCTTGCAATGAGTTGACrUCTGAG/iSpC3/ |
| SCRIB | 1 | fw | TAATACGACTCACTATAGGGAAATAGCAGCCAGGATGAAGTCATTGGAACArGGACGA/iSpC3/ |
| SCRIB | 1 | rev | TTCCAGGGACCTCAACTCCTCAGCAAArGTCCGT/iSpC3/ |
| SCRIB | 2 | fw | TAATACGACTCACTATAGGGAAATAACGCCTGTCACCGGACTTTGCTrGAGGAT/iSpC3/ |
| SCRIB | 2 | rev | TAGTTAGTCACAGGCCAGAACTCCTGTGrGGGTCC/iSpC3/ |
| FGFR2 | 2C | fw | TAATACGACTCACTATAGGGAAATAGGTGAATGTCACAGATGCCATCTCATCCrGGAGAC/iSpC3/ |
| FGFR2 | 2C | rev | GCTGGCAGAACTGTCAACCATGCAGArGTGAAG/iSpC3/ |
| FGFR2 | 2B | fw | TAATACGACTCACTATAGGGAAATAAAACAGCAAGCGCCTGGAAGAGAAArAGGAGG/iSpC3/ |
| FGFR2 | 2B | rev | GGCTGAGGTCCAAGTATTCCTCATTGGTrUGTGAT/iSpC3/ |
| FGFR1 | Alpha | fw | TAATACGACTCACTATAGGGAAATAGCAGTGACACCACCTACTTCTCCGTCrAATGTC/iSpC3/ |
| FGFR1 | Alpha | rev | GCTTCCAGAACGGTCAACCATGCAGArGTGATT/iSpC3/ |
| FGFR1 | Beta | fw | TAATACGACTCACTATAGGGAAATAGCACAAGCCACGGCGGACTCTrCCCGAT/iSpC3/ |

| | | | |
|---|---|---|---|
| FGFR1 | Beta | rev | GCTACGGGCATACGGTTTGGTTTGGrUGTTAC/iSpC3/ |
| PPFIBP1 | 1 | fw | TAATACGACTCACTATAGGGAAATAGAAGATCTTCGACAGTGCCTGAACAGGTrACAAGG/iSpC3/ |
| PPFIBP1 | 1 | rev | CAGAAGTTTCCATGCTTACAGTTGCTGGCrAATAAC/iSpC3/ |
| PPFIBP1 | 2 | fw | TAATACGACTCACTATAGGGAAATAGTCAAATGACAAATGGACACCTACCAGGGrAACGGG/iSpC3/ |
| PPFIBP1 | 2 | rev | CTGCTTTTCATACTGCATCTGACCCACCrUTGATC/iSpC3/ |
| PPFIBP1 | 3 | fw | TAATACGACTCACTATAGGGAAATACTCTCCACCCCCCAGTATAAAAGAACGTrGTGGAC/iSpC3/ |
| PPFIBP1 | 3 | rev | TGCTGCAGCATTTCTTCTGTGGCArUTCACA/iSpC3/ |
| PPFIBP1 | 4 | fw | TAATACGACTCACTATAGGGAAATAAAGTTCAGAGACACAGAGGGGCTGATTCrAGGAGG/iSpC3/ |
| PPFIBP1 | 4 | rev | CCGATCCTTTTCTTCATTTGCTGCCATCArAGGACC/iSpC3/ |
| SMAD2 | 1 | fw | TAATACGACTCACTATAGGGAAATAAGTGTTTTCAGTTCCGCCTCCAATCGrCCCATC/iSpC3/ |
| SMAD2 | 1 | rev | TTTCTTCCTGCCCATTCTGCTCTCCTrCCGCCC/iSpC3/ |
| SMAD2 | 2 | fw | TAATACGACTCACTATAGGGAAATACGGGAGGTTCGATACAAGAGGCTGTTTTrCCTAGA/iSpC3/ |
| SMAD2 | 2 | rev | AAGTTCTGTTAGGATCTCGGTGTGTCGGrGGCACC/iSpC3/ |
| DKK3 | 1A | fw | TAATACGACTCACTATAGGGAAATATTGCGGGCTCCCTCGGGTACrCGGCGA/iSpC3/ |
| DKK3 | 1A | rev | CGACTGGACCGAGTTGCGCTGrCGGGCA/iSpC3/ |
| DKK3 | 1B | fw | TAATACGACTCACTATAGGGAAATACTGAGCTCAGCCTCTCTTGGTGGATGrUGGGGA/iSpC3/ |
| DKK3 | 1B | rev | CGAACCCGGATCCTCTACGCTAATAGCrUCCCAG/iSpC3/ |
| CD44 | Standard | fw | TAATACGACTCACTATAGGGAAATATATTGTTAACCGTGATGGCACCCGCTrATGTCA/iSpC3/ |
| CD44 | Standard | rev | GATTCAGATCCATGAGTGGTATGGGACCCrCCCACC/iSpC3/ |

| | | | |
|---|---|---|---|
| CD44 | Variant | fw | TAATACGACTCACTATAGGGAAATAACCGACAGCACAGACAGAATCCCTrGCTACA/iSpC3/ |
| CD44 | Variant | rev | ATTTGAATGGCTTGGGTTCCACTGGGTCrCAGTCA/iSpC3/ |
| EXOC7 | Epithelial | fw | TAATACGACTCACTATAGGGAAATAGCATGATTTCCGAGTTAAGCACCTGTCCrGAGGCA/iSpC3/ |
| EXOC7 | Epithelial | fw | TAATACGACTCACTATAGGGAAATAAGTTAAGCACCTGTCCGAGGCCTTGAArCGACAG/iSpC3/ |
| EXOC7 | Epithelial | rev | AAACTCAGGCTTGGTCTGCTTGAGGTGTrCGCAGT/iSpC3/ |
| EXOC7 | mesenchymal | fw | TAATACGACTCACTATAGGGAAATATGATCAGTGGTGACGATGATCTGGAGGrCCCAGT/iSpC3/ |
| EXOC7 | Mesenchymal | rev | ATGTCATCTCTCCCTGGCCGCTTGArCTGGCC/iSpC3/ |
| FLNB | Hinge | fw | TAATACGACTCACTATAGGGAAATATGGAGGAGGCACCGGTAAATGCATGTrCCCCCC/iSpC3/ |
| FLNB | Hinge | rev | TCAGGGATGTGGCTGCCCATGTATTTrGATGTT/iSpC3/ |
| FLNB | No Hinge | fw | TAATACGACTCACTATAGGGAAATAGCAAGGTAGCCATCCTCAGAAGGTCAAArGTGTTC/iSpC3/ |
| FLNB | No Hinge | rev | ACGGCTGTGACTTCCCCATCTGTrGGCCAC/iSpC3/ |
| FLNB | No Hinge | rev | ATAGGCCTCTTCGGTCACCATGACAGTGArAGGGGA/iSpC3/ |

**Supplementary Table 7:** Isoform panel rhPCR primers.

### Supplementary Note 1

To better infer concentration values from fluorescence data, we mathematically modeled the Cas13 detection reaction with a system of differential equations. The model accounts for four primary processes: target transcription, Cas13-crRNA complexing, *cis*-cleavage, and *trans*-cleavage. While the complete reaction is certainly more complex, we find that this simplified description of the overall reaction is capable of modeling a wide range of experimentally generated fluorescence curves and inferring concentration values that correlate well with gold-standard methods.

$$DNA + T7 \xrightarrow{k_{cat}^{T7}} DNA + T7 + Target$$

$$C13 + crRNA \underset{k_r^{pre}}{\overset{k_f^{pre}}{\rightleftharpoons}} C13\text{-}crRNA$$

$$C13\text{-}crRNA + T \underset{k_r^{cis}}{\overset{k_f^{cis}}{\rightleftharpoons}} C13\text{-}crRNA\text{-}T \xrightarrow{k_{cat}^{cis}} AE$$

$$AE + FR \underset{k_r^{trans}}{\overset{k_f^{trans}}{\rightleftharpoons}} AE\text{-}FR \xrightarrow{k_{cat}^{trans}} AE + F$$

Key assumptions made in this model:

- The total amount of Cas13 enzyme is conserved
- The combined amount of quenched and unquenched fluorophore is conserved
- Transcription of target molecules does not follow Michaelis-Menten kinetics
- Cleaved reporter, active enzyme, and transcribed target are not present at the start of the reaction

### Supplementary Note 2

The reaction is modeled with a system of four ordinary differential equations describing changes in concentration of cleaved reporter (F), active enzyme (AE), target (T), and un-complexed Cas13 enzyme (C13).

$$\frac{dF}{dt} = \frac{k_{cat}^{trans}}{K_M^{trans}} \cdot AE \cdot FR$$

$$\frac{dAE}{dt} = \frac{k_{cat}^{cis}}{K_M^{cis}} \cdot C13\text{-}crRNA \cdot T$$

$$\frac{dT}{dt} = k_{cat}^{T7} \cdot T7_0 \cdot DNA + C13\text{-}crRNA \cdot T \cdot \left(\frac{k_r^{cis}}{K_M^{cis}} - k_f^{cis}\right) + AE \cdot T \cdot \left(\frac{k_r^{trans}}{K_M^{trans}} - k_f^{trans}\right)$$

$$\frac{dC13}{dt} = k_r^{pre} \cdot C13\text{-}crRNA - k_f^{pre} \cdot C13 \cdot crRNA$$

The following equations are used to enforce conservation:

$$E_{tot} = C13 + C13\text{-}crRNA + C13\text{-}crRNA\text{-}T + AE + AE\text{-}T + AE\text{-}FR$$

$$Rep_0 = FR + F + AE\text{-}FR$$

**Abbreviations**

| | |
|---|---|
| FR | Uncleaved reporter |
| $T7_0$ | T7 RNA polymerase |
| DNA | PCR product input |
| $E_{tot}$ | Initial Cas13 enzyme concentration |
| $Rep_0$ | Initial uncleaved reporter concentration |
